# A conserved mechanism for the retrieval of polyubiquitinated proteins from cilia

**DOI:** 10.1101/2025.04.24.650332

**Authors:** Sven M. Lange, Robyn J. Eisert, Alan Brown

**Author notes:** Correspondence should be addressed to A.B.

## Abstract

The temporospatial distribution of proteins within cilia is regulated by intraflagellar transport (IFT), wherein molecular trains shuttle between the cell body and cilium. Defects in this process impair various signal-transduction pathways and cause ciliopathies. Although K63-linked ubiquitination appears to trigger protein export from cilia, the mechanisms coupling polyubiquitinated proteins to IFT remain unclear. Using a multidisciplinary approach, we demonstrate that a complex of CFAP36, a conserved ciliary protein of previously unknown function, and ARL3, a GTPase involved in ciliary import, binds polyubiquitinated proteins and links them to retrograde IFT trains. CFAP36 uses a coincidence detection mechanism to simultaneously bind two IFT subunits accessible only in retrograde trains. Depleting CFAP36 accumulates K63-linked ubiquitin in cilia and disrupts Hedgehog signaling, a pathway reliant on the retrieval of ubiquitinated receptors. These findings advance our understanding of ubiquitin-mediated protein transport and ciliary homeostasis, and demonstrate how structural changes in IFT trains achieve cargo selectivity.

## Introduction

Cilia are organelles that extend from the cell surface and serve critical functions in various cellular processes including sensory perception, intercellular signaling, cell locomotion and fluid flow generation (Mitchison & Valente, 2017). Proper maintenance of the ciliary proteome, including the removal of damaged and mislocalized proteins as well as achieving the correct stoichiometry of signaling components, is fundamental for these processes. Imbalances in this maintenance are associated with a range of human ciliopathies including polycystic kidney disease, retinal degeneration, and developmental disorders such as Bardet-Biedl syndrome and Joubert syndrome (Moran *et al*., 2024).

The ciliary proteome is gated by the transition zone, a selective barrier at the ciliary base that separates the ciliary matrix and membrane from the remaining cell (Mercey *et al*., 2024). Many proteins are actively transported through this barrier by intraflagellar transport (IFT). The IFT machinery consists of two multiprotein subcomplexes, IFT-A and IFT-B, which assemble with kinesin and dynein motor proteins into megadalton-sized repeat structures referred to as IFT trains. These trains exist in two structurally distinct configurations to facilitate either anterograde (base-to-tip) or retrograde (tip-to-base) movement (Lacey *et al*., 2024, 2023; Pigino *et al*., 2009). Protein coupling to IFT trains often involves cargo adapters(Lechtreck, 2022) but the specific mechanisms by which they recognize cargo, associate with IFT trains, or differentiate between anterograde and retrograde transport, remain unknown.

Protein export from cilia has been proposed to be triggered by polyubiquitination, the covalent assembly of ubiquitin chains on target proteins. Early evidence for the role of ubiquitin in this process emerged from studies with *Chlamydomonas reinhardtii*, which demonstrate that their flagella (specialized motile cilia) contain ubiquitin, the machinery for ubiquitination, and polyubiquitinated signaling proteins (Huang *et al*., 2009). Mutant strains of *C. reinhardtii* with defective retrograde IFT show an increase in polyubiquitinated flagellar proteins (Huang *et al*., 2009), implicating IFT in the removal of polyubiquitinated proteins from flagella.

In mammalian primary cilia, polyubiquitination, specifically through K63-linked chains (Shinde *et al*., 2020), regulates signal transduction pathways by modulating receptor levels. For example, ubiquitination maintains Smoothened, a transmembrane protein of the Hedgehog (Hh) pathway, at low basal levels in cilia until pathway activation. Removing the ubiquitin moieties with a cilia-targeted deubiquitinase (Shinde *et al*., 2020; Lv *et al*., 2021) or deleting the ubiquitin-conjugating residues (Desai *et al*., 2020) causes Smoothened to accumulate in cilia even without Hh pathway activation, highlighting the essential role of ubiquitination in regulating this critical developmental and homeostatic pathway.

The IFT machinery is not thought to bind ubiquitin directly, suggesting the existence of protein(s) that couple polyubiquitinated proteins to IFT trains. A cargo adaptor for polyubiquitinated proteins would need to recognize ubiquitin chains assembled on different proteins, interact specifically with retrograde IFT trains, and maintain these interactions for the duration of transport towards the ciliary base. Current models propose that TOM1L2, a ubiquitin reader that functions in the endosomal sorting complexes required for transport (ESCRT) pathway (Shiba *et al*., 2004; Puertollano, 2005), a broadly utilized machinery not specific to cilia, is this adaptor. Evidence comes from proximity labelling studies localizing TOM1L2 within primary cilia (Mick *et al*., 2015), specific binding of TOM1L2 to K63-linked chains, and the accumulation of K63-linked ubiquitin chains and Smoothened in cilia following its genetic disruption (Shinde *et al*., 2023). RNAi-mediated knockdown of TOM1L2 also decreases the ciliary exit of two other ciliary transmembrane receptors, GPR161 and SSTR3 (Shinde *et al*., 2023), consistent with TOM1L2 being able to traffic diverse cargos. Based on *in vitro* binding assays, TOM1L2 is thought to interact with the IFT machinery through the BBSome (Shinde *et al*., 2023), an octameric complex that undergoes both anterograde and retrograde transport (Jin *et al*., 2010; Lechtreck *et al*., 2009). However, there is no direct evidence that the TOM1L2 protein itself can undergo IFT, raising questions about its placement in the ciliary protein retrieval pathway. It has also been conceptually perplexing that a pathway as specific to cilia as IFT would solely rely on the moonlighting function of a general ESCRT subunit for the critical function of coupling polyubiquitinated proteins to retrograde IFT trains. These limitations highlight a gap in our understanding of ciliary ubiquitin-mediated transport mechanisms.

Given the importance of protein export to ciliary homeostasis, we revisited the question of how polyubiquitinated proteins are recognized and retrieved from cilia. We started with the assumption that any dedicated, cilia-specific adaptor protein for polyubiquitinated proteins would have co-evolved with IFT and be present throughout ciliates. Using an unbiased proteomic screen of ubiquitin readers in flagellar extracts from two evolutionarily distant model organisms, we identified CFAP36 (also known as CCDC104 and BARTL1) as a highly conserved ciliary ubiquitin binder. We discovered that ubiquitin binding by CFAP36 is enhanced by ARL3, a small GTPase known previously only for its role in ciliary cargo unloading. We demonstrated that depletion of CFAP36 in mammalian cells results in the accumulation of K63-linked ubiquitin in cilia and impaired Hh signaling by disrupting the ciliary export of Hh signaling proteins. Through live-cell imaging, we showed that CFAP36 specifically undergoes retrograde IFT and identified that this is achieved through simultaneous detection of two IFT subunits that allows CFAP36 to selectively bind retrograde trains. Collectively, our findings identify a conserved, cilia-specific and ubiquitin-dependent mechanism for protein retrieval that is fundamental to ciliary function.

## Results

### CFAP36 is a cilia/flagella-specific polyubiquitin reader

A prior effort to identify ciliary ubiquitin readers used bovine retinal extracts as a source material (Shinde *et al*., 2023). Although retinal extracts are enriched with ciliary proteins, they also contain non-ciliary proteins from the cellular lysate that could potentially outcompete ciliary proteins for binding to polyubiquitin chains. To overcome this limitation, we used two unicellular organisms, *Chlamydomonas reinhardtii* and *Leishmania tarentolae*, from which we isolated highly pure flagella. Extracts from the purified flagella were passed over streptavidin beads coated with biotinylated K48- or K63-linked polyubiquitin chains, which are the most-abundant linkage types found in human cells (Swatek *et al*., 2019) (**Fig. 1A and Fig. S1A**). K48-linked ubiquitin chains typically target substrates for proteasomal degradation, whereas K63-linked ubiquitin chains are implicated in ciliary trafficking, endocytosis and protein complex regulation. Therefore, K63-linked chains are anticipated to be the preferred signal recognized by a ciliary ubiquitin adaptor. Proteins retained on the beads (**Fig. S1B-C**) were detected using quantitative multiplexed mass spectrometry through tandem mass tagging (TMT-MS), allowing for precise quantification of protein abundance across the different samples.

**Figure 1.**
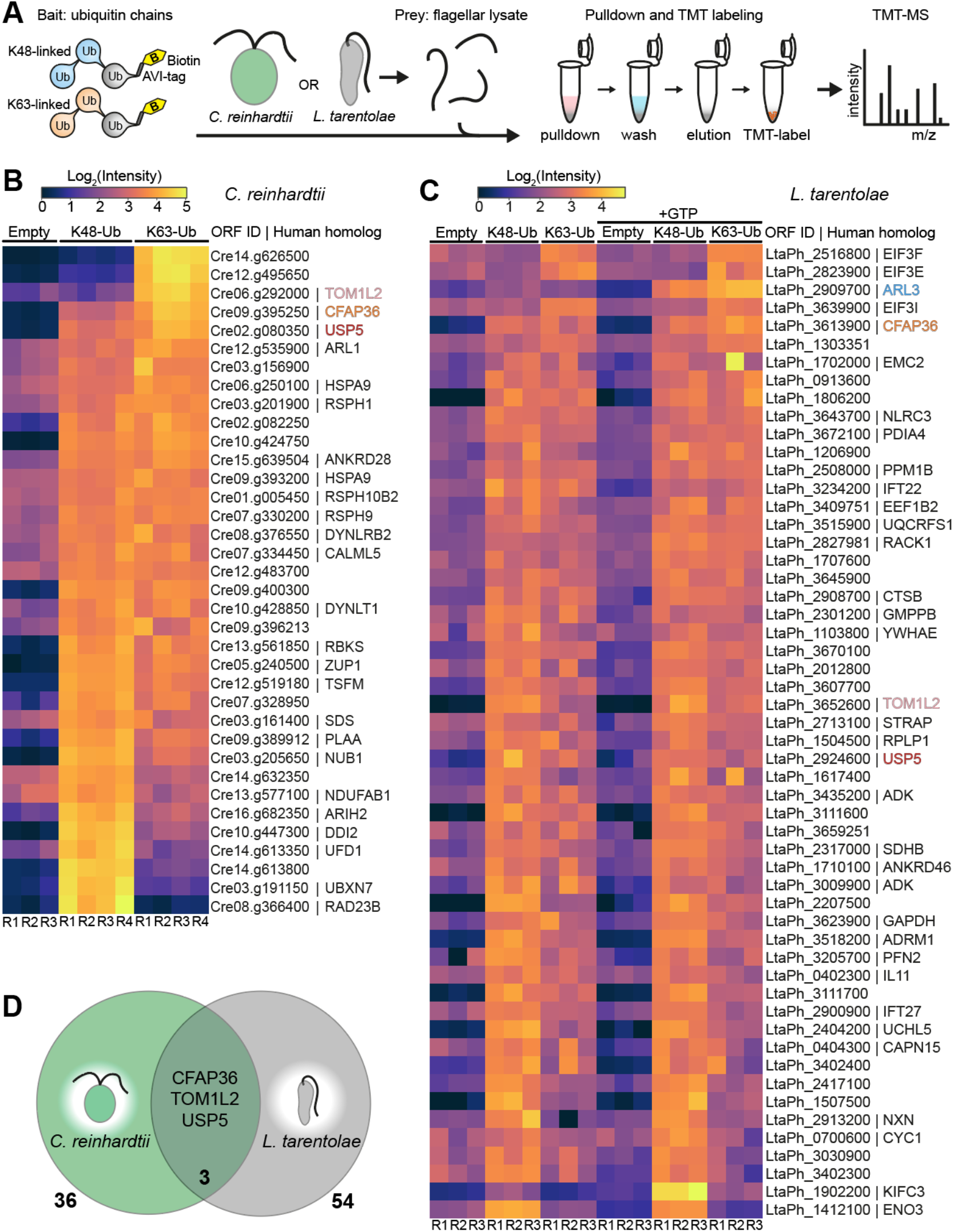
Identification of cilia-specific ubiquitin readers. (A) Schematic workflow of pulldown experiment from flagellar lysates using immobilized K48- and K63-linked ubiquitin chains and subsequent quantitative multiplexed mass spectrometry through tandem mass tagging (TMT-MS) analysis. (B, C) Heatmaps showing quantification of significantly enriched ubiquitin readers in *C. reinhardtii* (B) and *L. tarentolae* (C) flagellar lysates identified in triplicate or quadruplicate pulldowns (R1-R4). A one-way ANOVA was performed for each protein across experimental groups, followed by Ben-jamini-Hochberg false discovery rate (FDR) correction for multiple testing. Proteins with q-values below 0.01 were considered statistically significant, and those exhibiting a fold change greater than 2 compared to the empty beads control group were retained. (D) Venn diagram of proteins identified in the TMT-MS analysis of *C. reinhardtii* and *L. tarentolae* ubiquitin pulldowns. Only homologs of CFAP36, TOM1L2 and USP5 were identified in pulldowns from both organisms.

From *C. reinhardtii* flagella, 36 proteins exhibited enriched binding to the K48- or K63-linked ubiquitin chains (**Fig. 1B and Fig. S1D**), while 54 *L. tarentolae* proteins showed similar enrichment (**Fig. 1C and Fig. S1E**). Since *C. reinhardtii* (a green alga) and *L. tarentolae* (a protozoan parasite) belong to distinct evolutionary lineages (**Fig. S2A**), we rationalized that proteins present in both pulldowns could potentially correspond to a conserved ubiquitin reader that functions as an adaptor for the IFT machinery. Notably, only three proteins were detected in both samples: CFAP36, TOM1L2, and USP5 (**Fig. 1D**). We discounted USP5 as an IFT adaptor because of its well-characterized function as a deubiquitinase (Wilkinson *et al*., 1995) is incompatible with the need of an adaptor to make sustained interactions with polyubiquitinated proteins during transport. Given that TOM1L2 homologs have already been implicated in the export of polyubiquitinated proteins from cilia (Shinde *et al*., 2023), we focused instead on CFAP36, which bound both K48- and K63-linked polyubiquitin chains. CFAP36 is a highly conserved protein of unknown function with a phylogenetic distribution restricted to ciliated organisms (**Fig. S2A**) and a tissue-expression profile in humans that clusters with known ciliated cell types (Uhlén *et al*., 2015). Multiple proteomics studies have identified CFAP36 homologs in both motile and primary cilia (Pazour *et al*., 2005; May *et al*., 2021; McCafferty *et al*., 2024). These findings suggest that CFAP36 may play a conserved role in ciliary function.

### GTP-bound ARL3 enhances ubiquitin binding of CFAP36

Little is known about CFAP36 except that its N-terminal BART-like domain (**Fig. 2A**) forms a high-affinity complex with the GTP-bound form of the small GTPase ARL3 (Wright *et al*., 2011; Lokaj *et al*., 2015). ARL3 functions inside cilia as a cargo displacement factor for IFT, where it releases axonemal dynein precursors from anterograde IFT trains (Huang *et al*., 2024; Mali *et al*., 2024; Wang *et al*., 2025), and for lipidated intraflagellar transport (LIFT), where it displaces prenylated and myristoylated cargos from solubilizing factors PDE6D and UNC119 (Wright *et al*., 2011).

**Figure 2.**
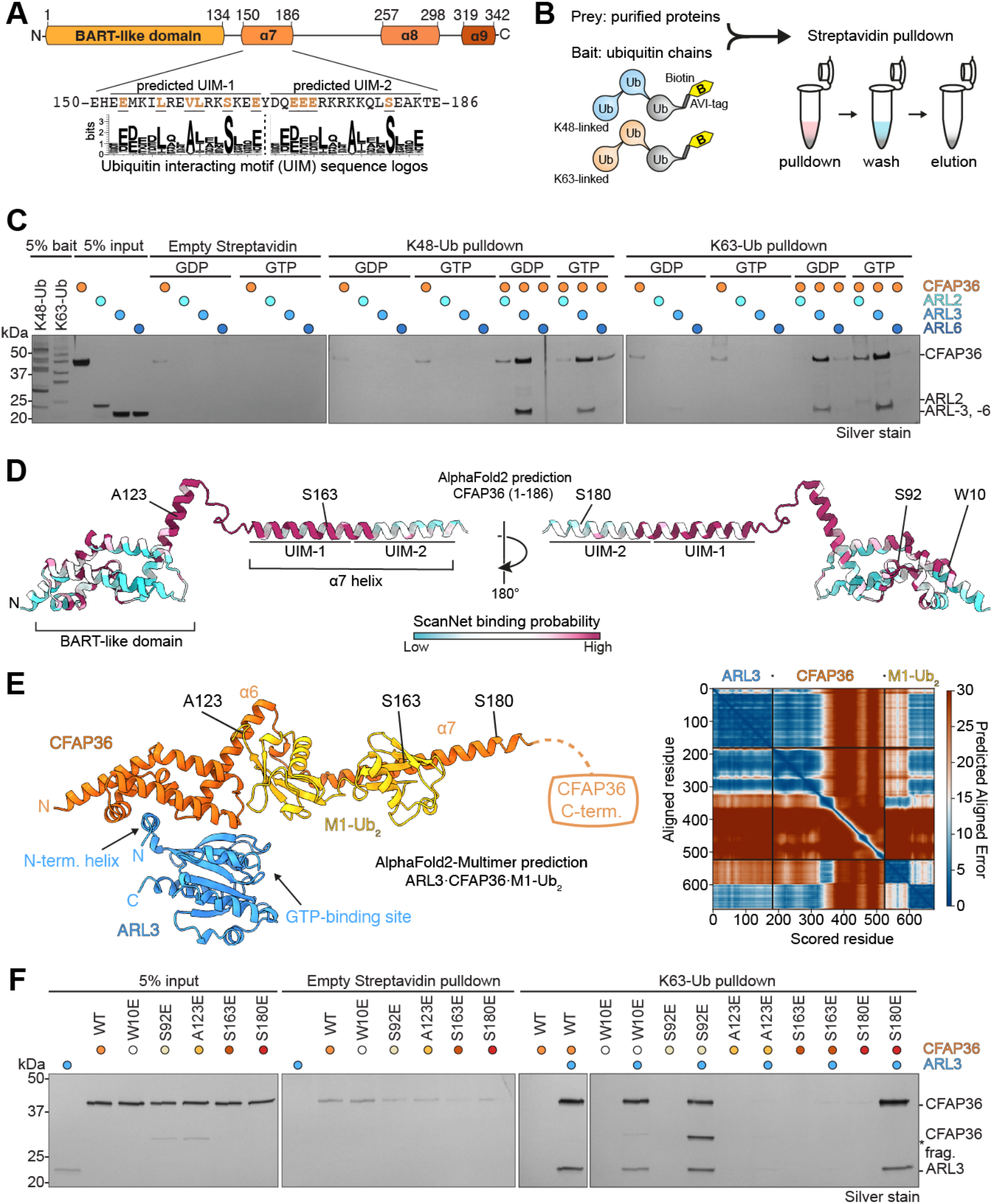
CFAP36 directly binds ubiquitin and ubiquitin binding is enhanced by small GTPase ARL3. (A)Domain overview of human CFAP36. CFAP36 has an N-terminal BART-like domain and three predicted C-terminal helices (α7, α8, and α9). Below, alignment of canonical ubiquitin-interacting motif (UIM) sequences to residues of the α7 helix identifies two potential UIMs. Residues matching the sequence motif are underlined and colored. (B)Schematic workflow of pulldown experiments between purified proteins and biotinylated ubiquitin chains shown in panels C and F. (C)Silver-stained SDS-PAGE analysis of the ability of recombinant, full-length porcine CFAP36 to bind streptavidin beads coated with K48- and K63-linked ubiquitin in the presence and absence of small GTPases, ARL2, ARL3 and ARL6 and GDP- or GTP-supplemented buffer. CFAP36 shows preferential binding to K48- and K63-linked ubiquitin in the presence of ARL3 and GTP. (D)AlphaFold2 model of human CFAP36 residues 1-186, colored by ScanNet binding site probability. Highlighted residues (with porcine numbering) are those targeted for mutational analysis shown in panel F. (E)AlphaFold2-Multimer model of a CFAP36•ARL3•diubiquitin complex with predicted aligned error (PAE) plot showing regions of high confidence (blue) and low confidence (red). Diubiquitin was modeled as tandem copies of ubiquitin (M1-linkage) because it is not yet possible to model other linkage types in either AlphaFold2 or AlphaFold3. Highlighted residues are those targeted for mutational analysis shown in panel F. (F)Silver-stained SDS-PAGE of K63-linked ubiquitin pulldown with recombinant porcine CFAP36 variants with mutations in predicted UIMs and regions of high protein binding probability. Substitution of residues in the α6 and α7 helices (A123E and S163E, respectively) completely abolish ubiquitin binding in vitro.

ARL3 was detected in our ubiquitin pulldown from *L. tarentolae* flagella following the addition of GTP (**Fig. 1C**) with similar K48/K63-ubiquitin binding preferences to CFAP36, suggesting that ARL3 and CFAP36 bind polyubiquitin chains as a complex *in vivo*. To further test this possibility, we purified recombinant porcine CFAP36 and performed *in vitro* binding experiments with K48- and K63-linked polyubiquitin chains in the presence of ARL3•GDP or ARL3•GTP (**Fig. 2B, C**). The results showed that both ARL3•GDP and ARL3•GTP enhanced the binding of CFAP36 for polyubiquitin chains, with greatest binding in the presence of GTP. ARL3 showed little binding to polyubiquitin chains on its own. Because the interaction between CFAP36 and ARL3 is largely mediated by the amphipathic N-terminal helix of ARL3 (Lokaj *et al*., 2015), we included two related small GTPases, ARL2 and ARL6, that contain similar amphipathic N-terminal helices as controls. ARL2, a close non-ciliary homolog of ARL3 that binds CFAP36 with 10-fold lower affinity (Lokaj *et al*., 2015), also enhanced binding between CFAP36 and ubiquitin in the GTP-bound state (**Fig. 2C**). ARL6, a more distantly related GTPase that binds the BBSome(Jin *et al*., 2010), had no effect (**Fig. 2C**). These data demonstrate that CFAP36 can bind polyubiquitin chains and that its interaction is enhanced by GTP-bound ARL3.

### CFAP36 interacts with ubiquitin through a non-canonical ubiquitin-interacting motif

To elucidate the molecular basis for the interaction between CFAP36 with ubiquitin, we analyzed its sequence and identified two candidate non-canonical ubiquitin-interacting motifs (UIMs) within helix α7, which follows the BART domain (**Fig. 2A**). A UIM is a short α-helical element characterized by a conserved serine residue positioned among acidic and hydrophobic residues, which binds to the hydrophobic I44-patch of a single ubiquitin (Young *et al*., 1998; Hofmann & Falquet, 2001) but can confer linkage specificity when occurring in tandem (Sato *et al*., 2009). The first potential UIM (UIM-1) is closer in sequence to the consensus for a UIM and has a higher probability of being a protein binding interface than UIM-2 based on an analysis using ScanNet (**Fig. 2D**), a deep-learning tool designed to predict protein-protein interfaces (Tubiana *et al*., 2022). UIM-1 is also predicted to interact with ubiquitin in an AlphaFold2-Multimer model of the ternary complex of CFAP36, ARL3 and a diubiquitin (**Fig. 2E**). AlphaFold2-Multimer also predicted a second interface between ubiquitin and the α6 helix of the BART domain, which does not contain a UIM.

To experimentally test these predicted interfaces *in vitro*, we generated five mutant proteins, each containing a single amino acid substituted with a glutamate. The substituted residues included S163 in UIM-1, S180 in UIM-2, A123 of helix α6, and two additional residues in the BART domain (W10 and S92) located in surface patches with high ScanNet protein binding probabilities but no predicted ubiquitin binding (**Fig. 2D**). We tested this set of mutant proteins for their ability to bind K63-linked ubiquitin chains in the presence and absence of ARL3•GTP. The binding assays revealed that the W10E and S92E mutations within the BART domain, as well as the S180E substitution in UIM-2, had no effect on the ability of CFAP36 to bind ubiquitin (**Fig. 2F**). UIM-2 is therefore not necessary for ubiquitin binding. In contrast, the S163E substitution in UIM-1 and the A123E substitution in helix α6 completely abrogated binding. These findings indicate that both the α6 and α7 helices are essential for CFAP36 to bind ubiquitin.

Next, we speculated that ARL3•GTP might enhance the affinity of CFAP36 for ubiquitin chains by alleviating autoinhibition. We explored two possibilities. First, we considered the existence of an autoinhibited homodimeric form of CFAP36, as predicted by a high-throughput AlphaFold2 screen (Schweke *et al*., 2024). However, size exclusion chromatography coupled with multi-angle light scattering (SEC-MALS) clearly demonstrated that purified recombinant full-length CFAP36 was monomeric (**Fig. S3A**). Second, we investigated whether the C-terminal α8 and α9 helices of CFAP36 might interfere with ubiquitin binding. To test this, we generated a C-terminal truncation of CFAP36 (retaining residues 1-186, including the BART-like domain and helix α7) and examined its ability to bind K63-linked ubiquitin chains in the absence and presence of ARL3•GTP (**Fig. S3B**). Our findings revealed that while removal of the C-terminal helices enhanced CFAP36 binding to K63-linked ubiquitin chains, the binding levels remained lower than in the presence of ARL3•GTP. These results suggest that while the C-terminal helices may lower the affinity of CFAP36 for ubiquitin, ARL3•GTP likely promotes binding through additional mechanisms.

### CFAP36 has broad ubiquitin-dependent specificity for membrane-associated proteins

The identification of a single amino acid substitution (S163E) in CFAP36 that abolished ubiquitin binding (**Fig. 2F**) provided a tool to identify its ubiquitin-dependent binding partners. Using both wild-type (WT) and mutant (S163E) recombinant porcine CFAP36 proteins as bait, we performed pull-down studies with cilia-enriched extract from porcine respiratory tissue as a source of endogenously ubiquitinated ciliary proteins (**Fig. 3A**). As CFAP36 ubiquitin binding was enhanced when complexed with ARL3•GTP, we fused CFAP36 with ARL3 (ARL3-CFAP36, see Methods) and added GTP during the pulldown. Following incubation, proteins eluted from unbound beads and those coated with either WT or S163E mutant were analyzed using TMT-MS (**Fig. 3B**). We identified 49 proteins whose interaction with CFAP36 was lost in the ubiquitin-binding deficient mutant (**Fig. 3B**). One of these proteins was ubiquitin itself, validating our approach. Among the other identified proteins, five were E3 ubiquitin ligases, potentially reflecting their autoubiquitination. Thirty-five (71%) were integral membrane proteins including epidermal growth factor receptor (EGFR), transmembrane ATPases, and members of the solute carrier family, many of which are known targets of K63-linked polyubiquitination (Galcheva-Gargova *et al*., 1995; Huang *et al*., 2006; Argenzio *et al*., 2011). Thirty-seven (80%) had CilioGenics scores (Pir *et al*., 2024) of >0.5 indicating mild to high probability of being a ciliary protein. This finding suggested that the CFAP36•ARL3•GTP complex is capable of binding a diverse array of polyubiquitinated soluble and transmembrane proteins, supporting the hypothesis that it may function as a general adaptor for the ciliary export of polyubiquitinated proteins.

**Figure 3.**
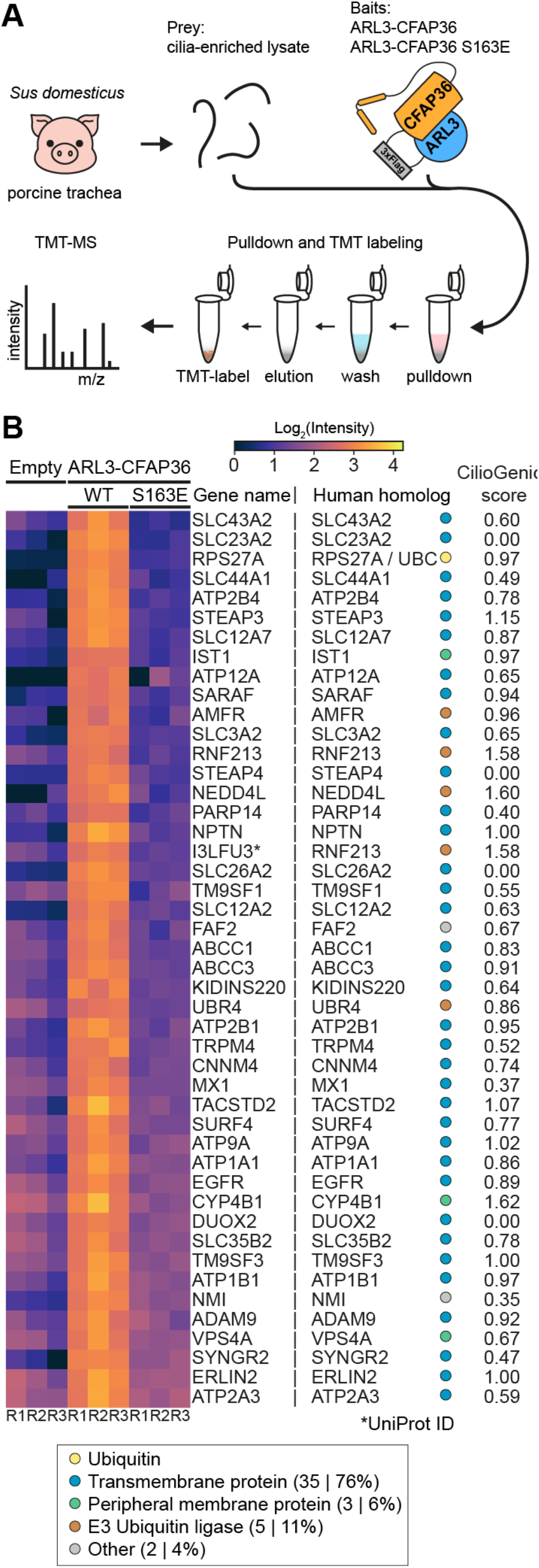
ARL3-CFAP36 fusion interacts with diverse membrane-associated proteins via ubiquitin. (A)Schematic workflow of the pulldown used to identify proteins within porcine trachea cilia extracts that interact with ARL3-CFAP36 fusion proteins in a ubiquitin-dependent manner. (B)Heatmap showing proteins significantly reduced following pulldown with ubiquitin-binding deficient ARL3-CFAP36 S163E mutant compared to wild type (WT). Each experiment was performed in triplicate (R1-R3). A one-way ANOVA was performed for each protein across experimental groups, followed by Benjamini-Hochberg FDR correction for multiple testing. Proteins with q-values below 0.01 were considered statistically significant, and those exhibiting a fold-change of the WT group greater than 2 compared to the empty beads control group and S163E group were retained. Identified proteins were classified as ubiquitin, trans-membrane, peripheral membrane-associated, E3 ubiquitin ligase, or other. 80% of the identified proteins have a CilioGenics score(Pir *et al*., 2024) of >0.5 and >1.0 indicating mild and high probability for cilia localization, respectively.

### CFAP36 depletion disrupts ubiquitin-mediated ciliary retrieval

We next investigated whether retrieval of ubiquitinated proteins from the cilium is impaired by CFAP36 loss. To address this, we performed CFAP36 knockdown (depletion) experiments in IMCD3 cells, a mouse kidney cell line characterized by ∼6 μm-long primary cilia (**Fig. 4A**). Using synthetic small interfering RNAs (siRNAs) targeting *Cfap36*, we achieved knockdown efficiencies of approximately 98% as determined by immunoblot analysis with a CFAP36 antibody (**Fig. 4B**). Knockdown of CFAP36 resulted in a significant increase in ciliary K63-linked ubiquitin compared to control cells treated with non-targeting siRNAs (**Fig. 4C**). Furthermore, we observed an accumulation of the Hh pathway transmembrane proteins Smoothened (**Fig. 4D**) and GPR161 (**Fig. 4E**) in cilia following CFAP36 depletion. These findings indicate that CFAP36 knockdown enhances both ciliary K63-ubiquitin levels and the retention of transmembrane receptors that rely on ubiquitination and IFT for their retrieval from cilia.

**Figure 4.**
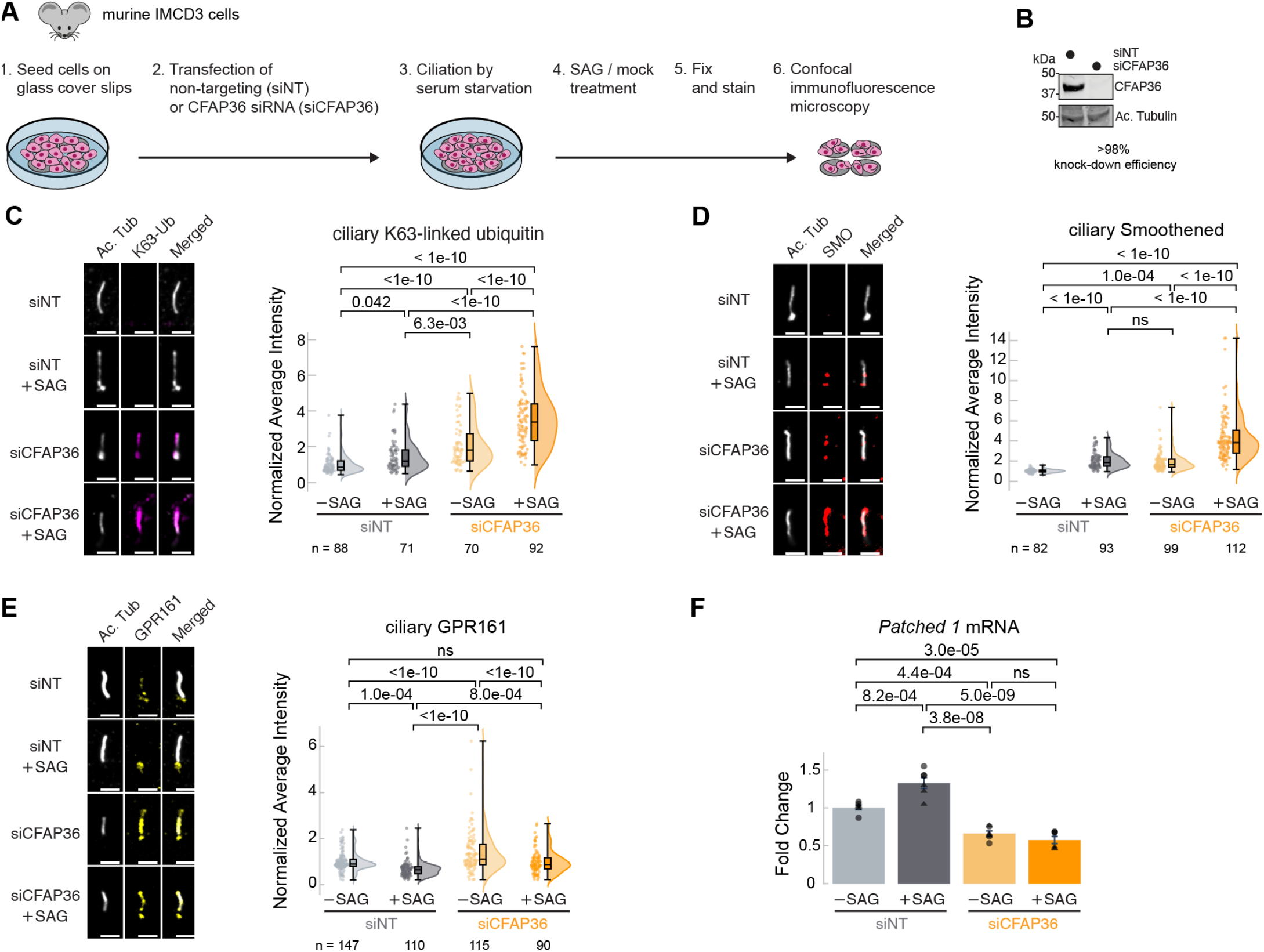
CFAP36 depletion leads to accumulation of K63-linked ubiquitin in cilia and disrupts ciliary retrieval of GPCRs. (A)Schematic showing sample preparation workflow for immunofluorescence microscopy. Ciliation in murine IMCD3 cells was induced by removing serum from the medium. (B)Immunoblot analysis of whole-cell lysates from ciliated IMCD3 cells transfected with non-targeting (siNT) or CFAP36-specific (siCFAP36) siRNA. The top panel shows detection of CFAP36 using an anti-CFAP36 antibody. The bottom panel shows detection of anti-acetylated tubulin (Ac. Tub) as a loading control. (C-E) Left: Representative images of cilia stained for ciliary marker acetylated tubulin (Ac. Tub) and either (C) K63-linked ubiquitin (K63-Ub), (D) Smoothened (SMO), or (E) G Protein-Coupled Receptor 161 (GPR161). Right: Raincloud plots of average cilia-localized signal intensities normalized to siNT control treatment. The bottom and top of the box represents the first (Q1) and third (Q3) quartiles, respectively. The median is marked by a line inside the box. The whiskers extend to the highest and the lowest measurements. The number of quantified cilia per condition (minimum = 70) are indicated below the graphs. Statistical significance of the differences was assessed using one-way ANOVA followed by Tukey’s Honest Significant Difference (HSD) test with a significance level of α = 0.05. Results are displayed as p-values. (F) Quantitative real-time polymerase chain reaction (qRT-PCR) analysis of *Patched 1* mRNA levels in IMCD3 cells transfected with siNT or siCFAP36, with and without SAG treatment. Data are expressed as relative expression levels of *Patched 1*, normalized against the reference gene RPL27. Statistical significance of the differences between groups of six replicates was assessed using One-way ANOVA followed by Tukey’s HSD test with a significance level of α = 0.05. Whiskers of bar graphs indicate standard deviations.

We also performed parallel experiments in the presence of 200 nM Smoothened Agonist (SAG), a small molecule activator of the Hh pathway known to elevate Smoothened levels in cilia (Rohatgi *et al*., 2007) while decreasing ciliary GPR161 levels (Mukhopadhyay *et al*., 2013). We observed this expected response to SAG stimulation in IMCD3 cells treated with both non-targeting and *Cfap36*-targeting siRNAs. However, CFAP36-deficient cells displayed a hyperaccumulation of Smoothened in cilia (**Fig. 4D**), while GPR161 levels did not decrease beyond baseline levels observed in unstimulated control cells (**Fig. 4E**). These results indicate that transmembrane receptors can enter but not exit cilia without CFAP36. Thus, CFAP36 depletion specifically impacts ciliary export and disrupts the normal ubiquitin-dependent redistribution of Hh signaling pathway components in response to SAG treatment.

The movement of proteins in and out of cilia is critical for eliciting a transcriptional response to Hh signaling(Rohatgi *et al*., 2007). To assess the impact of transmembrane protein mislocalization in CFAP36-depleted cells on Hh signaling, we used quantitative real-time polymerase chain reaction (qRT-PCR) to measure the expression of *Patched1* (*Ptch1*), a target gene of the Hh signaling that acts as a negative regulator of the pathway (Taipale *et al*., 2002) (**Fig. 4F**). As expected, *Ptch1* expression increased in response to SAG-induced signaling in IMCD3 control cells, with a fold change similar to reported in IMCD3 cells (Li *et al*., 2011). In contrast, cells lacking CFAP36 showed no transcriptional response to SAG-induced signaling, with baseline *Ptch1* transcript levels remaining significantly lower than those in control cells where CFAP36 was present. These results demonstrate that CFAP36 is an essential component of the Hh signaling pathway and is required for appropriate gene expression following SAG stimulation.

### CFAP36 undergoes retrograde IFT

We next tested the model that CFAP36 functions as an IFT cargo adapter for ubiquitinated proteins by assessing whether it is capable of moving with IFT trains. To do this, we used total internal reflection fluorescence (TIRF) microscopy, which is particularly well suited to visualizing IFT in *C. reinhardtii* flagella (Engel *et al*., 2009). To be able to track CFAP36 movement, we used CRISPR/Cas9 gene editing to endogenously tag the C-terminus of CFAP36 with a fluorescent protein, mStayGold (Ivorra-Molla *et al*., 2024) (**Fig. S5A**), in a *C. reinhardtii* strain expressing mScarlet fused to the N-terminus of IFT54 (Wingfield *et al*., 2017), an integral component of the IFT-B subcomplex. In these double-tagged strains, CFAP36-mStayGold underwent retrograde transport and co-localized with mScarlet-IFT54, with few instances of anterograde transport (**Fig. 5A, B**). The prevalence of retrograde traces is particularly notable as this technique is biased toward the detection of anterograde transport because photobleaching depresses the retrograde signal. It further suggests that most CFAP36 diffuses passively into flagella before being actively transported, via retrograde transport, back to the cell body. Not all CFAP36-mStayGold traces have corresponding mScarlet-IFT54 traces; because there is no difference in the kinetics between the two types, we attribute this to reduced photobleaching of mStayGold relative to mScarlet rather than examples of CFAP36 moving independently of IFT.

**Figure 5.**
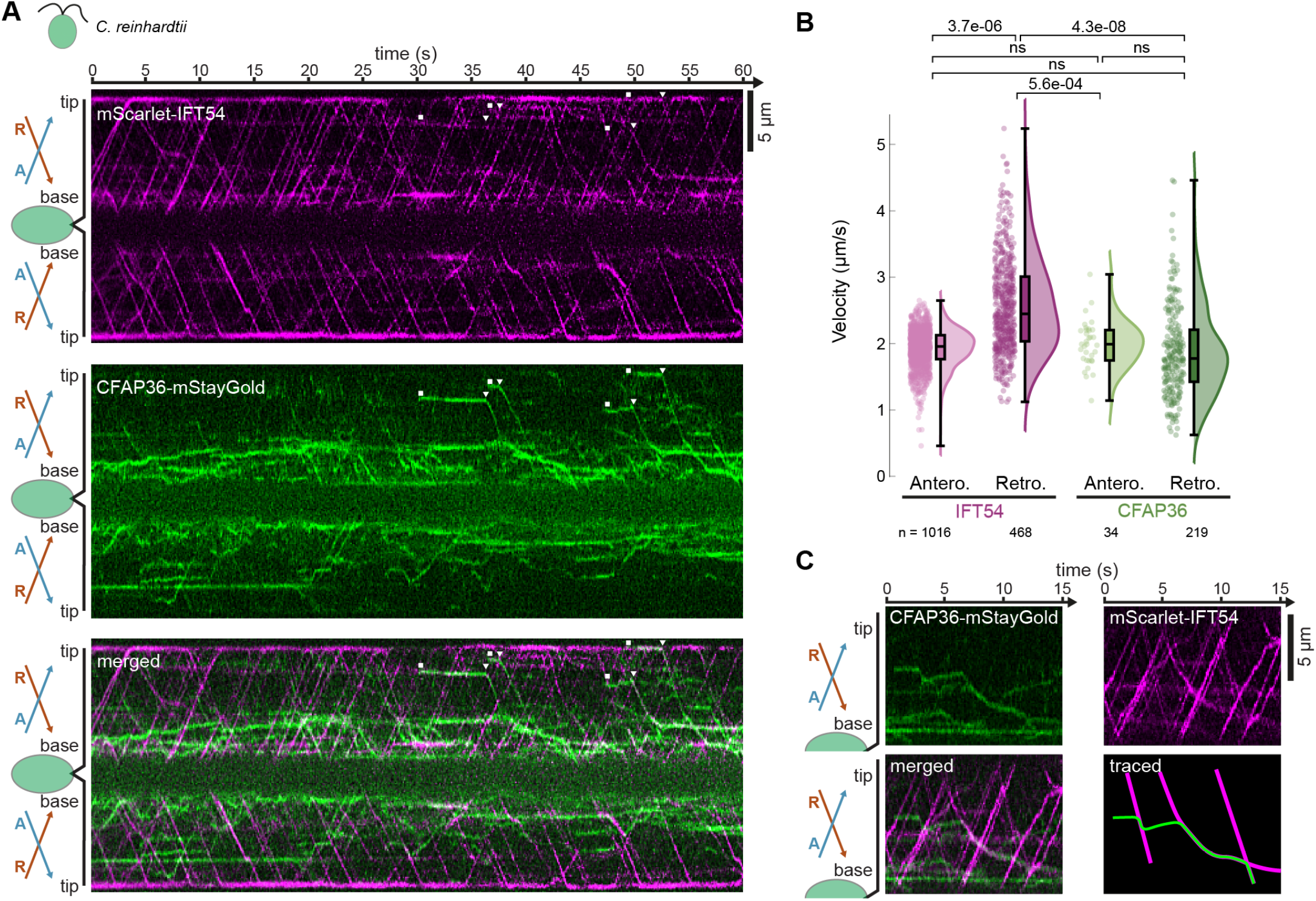
CFAP36 preferentially travels via retrograde IFT. (A)Representative kymographs of *C. reinhardtii* flagella showing 60 s time lapse of mScarlet-IFT54 (top) and CFAP36-mStayGold (middle) imaged by TIRF microscopy. The bottom panel represents a merge of both signals. White squares mark the appearance of selected stationary CFAP36 traces, whereas white triangles denote the start of retrograde transport. Schematics on the left indicate orientation of cell body and flagella with indicated anterograde (A) and retrograde (R) IFT movement directions. (B)Raincloud plots of velocities calculated from anterograde and retrograde traces of mScarlet-IFT54 and CFAP36-mStayGold, respectively. Each point represents an individual measurement. In the box plot, the central line represents the median, with the bottom and top of the box representing the first (Q1) and third (Q3) quartiles, respectively. The whiskers extend to the highest and lowest measurements. Statistical significance was determined using one-way ANOVA followed by Tukey’s HSD test with results displayed as p-values. Number of measurements are indicated below the graphs. (C)Section from a kymograph showing disjointed retrograde transport of CFAP36-mStayGold and decelerated IFT trains upon CFAP36 loading.

The CFAP36-mStayGold retrograde traces initiate from various locations within the flagellum including the proximal region, indicating that train recognition is not restricted to the flagellar tip. Many of the CFAP36-mStayGold traces also begin with a stationary phase that can last several seconds, during which changes in intensity may indicate clustering of CFAP36 in the flagellum as it awaits a retrograde IFT train. We also observed that CFAP36 can detach from trains prior to the flagellar base before becoming cargos for a second train (**Fig. 5C**), suggesting that the affinity of coupling may be relatively weak. Although TIRF shows clear progressive retrograde movement of CFAP36 toward the flagellar base, we are unable to reliably capture export from the flagellum because the base is at variable distances from the evanescent field created by TIRF illumination.

The median velocity of CFAP36-mStayGold during retrograde transport (1.8 μm s^-1^) was slower than the median velocity (2.4 μm s^-1^) of retrograde transport measured from mScarlet-IFT54 traces (**Fig. 5B**). The velocity of CFAP36-associated trains also appears to slow further as it approaches the flagellar base (**Fig. 5C**). In contrast, the few recorded instances of CFAP36-mStayGold anterograde transport (2.0 μm s^-1^) did not show a statistically significant difference in velocity from anterograde moving mScarlet-IFT54 (2.0 μm s^-1^). This suggests that retrograde IFT trains carrying CFAP36 have distinct kinetics from those that do not. Whether this is due to CFAP36 directly regulating the mobility of retrograde trains or CFAP36-associated cargos increasing drag remains to be determined. Collectively, these findings support the model that CFAP36 functions as an IFT cargo adapter, participating in the retrograde transport of ubiquitinated proteins.

### CFAP36 specifically recognizes retrograde trains

To address the question of how CFAP36 specifically interacts with retrograde but not anterograde IFT trains, we searched for potential binding partners of CFAP36 by performing an unbiased AlphaFold2-based prediction screen using a list of ∼3300 potential ciliary proteins compiled from five different sources (**Fig. 6A**). Each protein on the list was paired with CFAP36 and structures predicted with AlphaFold2-multimer, with the results analyzed using the Structure Prediction and Omics informed Classifier (SPOC), a machine-learning based approach developed to distinguish between true and false AlphaFold2-Multimer predictions (Schmid & Walter, 2025). The solutions were ranked by compact SPOC (cSPOC) score (**Fig. 6B**), with higher values more likely to represent *bona fide* interactions. The presence of ARL3 and UBC (polyubiquitin) among the top solutions confirmed the effectiveness of the approach for identifying known interactions.

**Figure 6.**
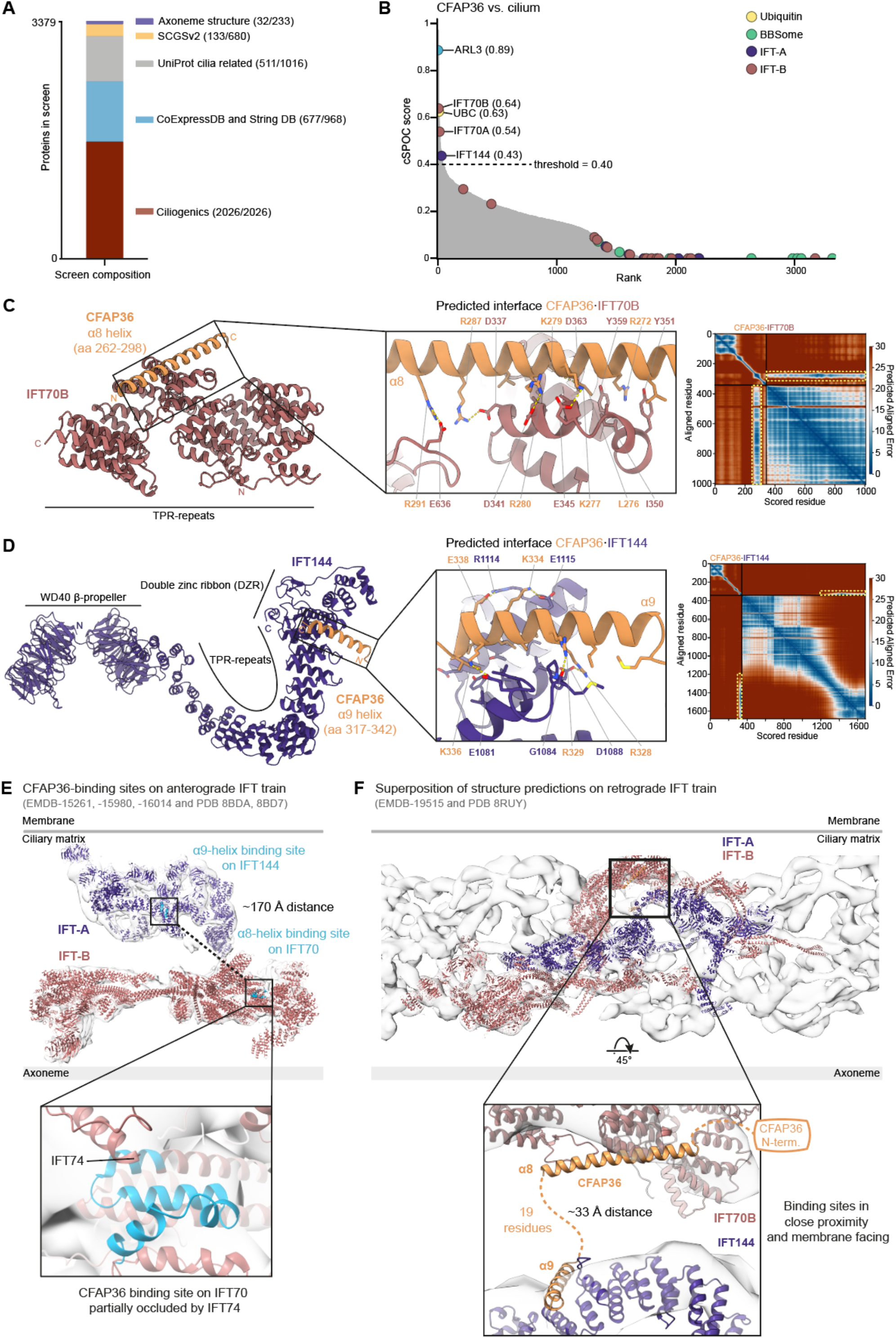
AlphaFold2 identifies two IFT subunits as CFAP36 interactors for coincidence detection of retrograde trains. (A)Cumulative composition of the human ciliary proteome from five sources. Numbers in parentheses indicate the unique entries added from each source. (B)Plot showing cSPOC(Schmid & Walter, 2025) score versus rank for pairwise AlphaFold2-Multimer predictions between CFAP36 and the human ciliary proteome. The threshold for identifying a true interaction (0.40) was set by the elbow point of the curve. Pairwise predictions involving CFAP36 and ubiquitin and subunits of the IFT-A, IFT-B and BBSome complexes are represented as colored dots. Those above the threshold are annotated with their protein name and cSPOC score. (C, D) Left, Predicted human protein complexes of IFT70B with CFAP36 helix α8 (C) or IFT144 with CFAP36 helix α9 (D), including a close-up view of the predicted interface residues. Right, Predicted aligned error (PAE) plots from AlphaFold2-Multimer, with interacting helices highlighted in yellow dashed boxes. (E)Cryo-ET-derived map and atomic model of the *C. reinhardtii* anterograde IFT train with IFT-A colored purple and IFT-B colored rust. Predicted CFAP36-binding sites on IFT70 and IFT144 are highlighted in light blue and distance indicated by dashed line. Zoomed view shows that the predicted α8-helix binding site is partially obscured by IFT74. (F)Cryo-ET-derived map and atomic model of the *C. reinhardtii* retrograde IFT train colored as in panel E. Zoomed-in view shows IFT70 and IFT144 with superimposed CFAP36 complex predictions. Dashed orange lines indicate gap between α8- and α9-helices, and connection to CFAP36 N-terminus.

Three IFT subunits also scored above the threshold for likely true interactors; IFT70 paralogs A and B and IFT144 (also known as WDR19). Notably, the cSPOC score for IFT70B is even higher than the score for polyubiquitin. Inspection of the interfaces revealed that IFT70A/B and IFT144 are predicted to engage different C-terminal helices of CFAP36: IFT70A/B with α8 (**Fig. 6C**), and IFT144 with α9 (**Fig. 6D**), suggesting that CFAP36 could be capable of binding both proteins simultaneously. In each case, the interaction involves tetratricopeptide repeats (TPRs) of the IFT subunits, which contribute both hydrophobic and electrostatic residues to the interface. The predicted alignment error (PAE) for these interactions is low, signifying a high confidence interaction (**Fig. 6C,D**). IFT70A and IFT70B are over 95% identical and the only difference in the CFAP36 binding site is a threonine to isoleucine substitution in IFT70B. This small change may explain why IFT70B scores slightly better than IFT70A. Reciprocal AlphaFold2-Multimer screens using IFT70 or IFT144 as the bait also identified CFAP36 among their top scoring binding partners (**Fig. S4**).

IFT70 and IFT144 belong to different subcomplexes of the IFT machinery: IFT-B and IFT-A, respectively. The orientation of these subcomplexes with respect to each other changes in the transition from anterograde to retrograde trains, as revealed by electron cryotomography (cryo-ET) of *C. reinhardtii* IFT trains (Lacey *et al*., 2024, 2023). In anterograde IFT trains, IFT70 and IFT144 are 170 Å apart (**Fig. 6E**) – a distance too far to allow simultaneous recognition by CFAP36. Furthermore, the binding site on IFT70 is partially obstructed by another subunit – IFT74 (**Fig. 6E**). In contrast, IFT70 and IFT144 are adjacent to one another in the retrograde train (**Fig. 6F**). The predicted binding sites are solvent exposed, membrane-facing, and the 33 Å distance between them could be easily spanned by the linker between the α8 and α9 helices (19 residues in humans). We therefore propose a coincidence detection mechanism that allows CFAP36 to specifically bind retrograde IFT trains by sensing the spatial organization of IFT-A and -B subcomplexes.

### Mutations at IFT-interaction interfaces disrupt CFAP36 retrograde transport

To test our proposed model of coincidence detection *in vivo*, we used CRISPR/Cas9 to introduce structure-guided mutations into the endogenous *Cfap36* locus of the *C. reinhardtii* IFT54-mScarlet strain (**Fig. S5B**). To disrupt the interaction between CFAP36 and IFT70, we modified the CFAP36 α8 helix by substituting a hydrophobic leucine residue (L438) and a salt-bridge-forming lysine residue (K442) with glutamate residues (**Fig. 7B**). To disrupt the interaction with IFT144, we modified the CFAP36 α9 helix by replacing an interfacial alanine residue (A493) with a bulkier glutamate and disrupted a salt bridge by replacing K495 with an alanine residue (**Fig. 7B**). We also generated a strain with all four mutations (L438E, K442E, A493E, K495A) to disrupt interactions involving both the α8 and α9 helices. TIRF microscopy analyses of these three strains revealed that while the mutations did not prevent CFAP36 from entering flagella, they inhibited its movement within the flagellum. instances of retrograde or anterograde movement of CFAP36 were rare (**Fig. 7D-F**). In contrast, IFT dynamics, monitored by IFT54-mScarlet, were unaffected (**Fig. S6A-F**). These findings demonstrate that interactions involving both the α8 and α9 helices are necessary for coupling CFAP36 to retrograde IFT trains.

**Figure 7.**
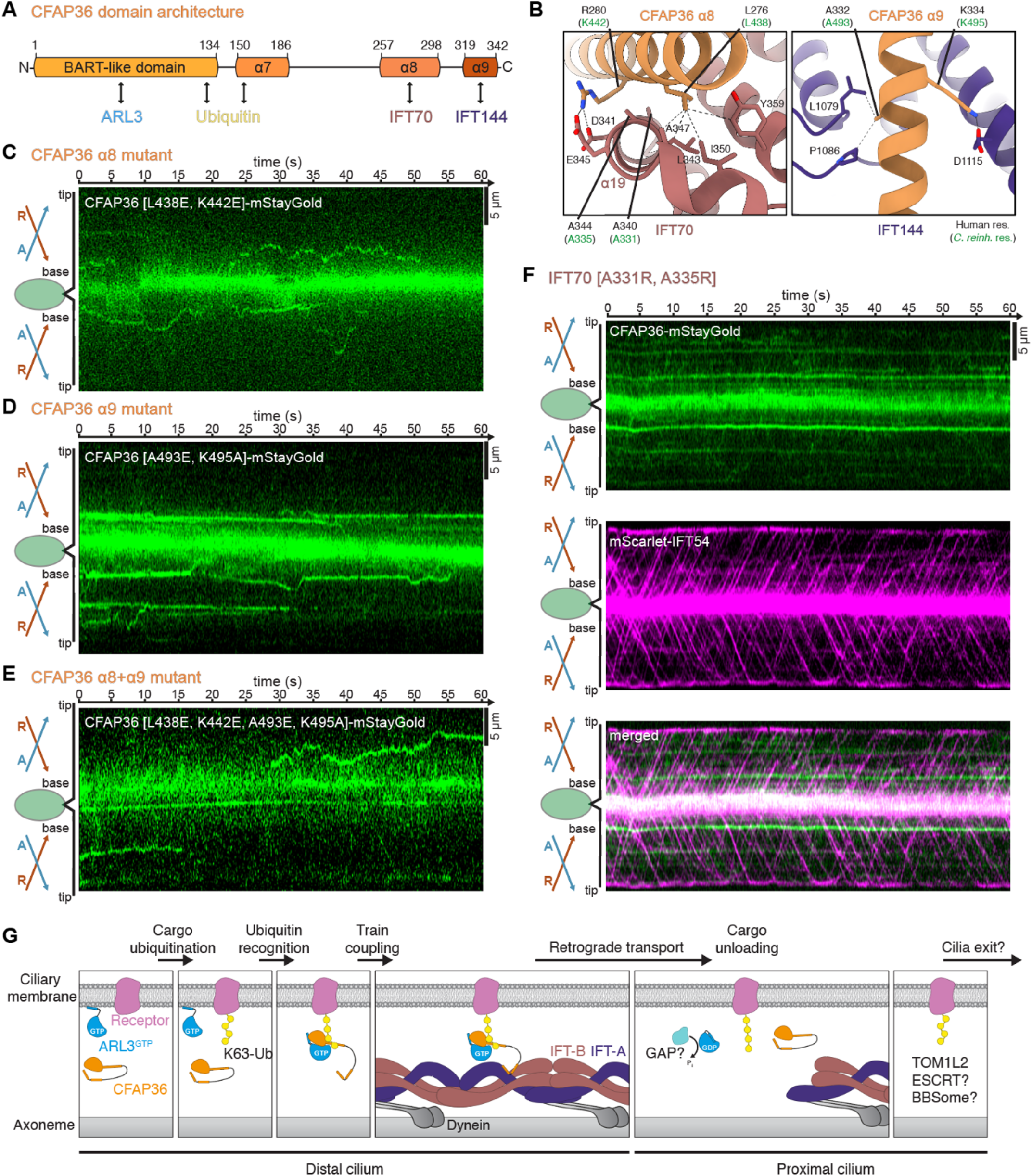
In vivo validation of CFAP36 coincidence detection of retrograde IFT. (A)Schematic domain overview and interaction landscape of CFAP36 (with human numbering). (B)Detailed view of residues selected for mutational analysis based on predicted human CFAP36•IFT complexes with corresponding residues in *C. reinhardtii* shown in parentheses in green. Dashed lines indicate electrostatic and hydrophobic interactions of selected amino acid side chains. (C-E) Representative kymographs showing 60 s time lapse of *C. reinhardtii* flagella with CFAP36-mStayGold harboring disruptive mutations in α8 (C), α9 (D) or both (E) interfaces imaged by TIRF microscopy. The strong central signal intensity is due to epi-fluorescence signal from the cell body. (F)Representative kymographs depicting a 60-second time lapse of *C. reinhardtii* flagella from the IFT70 [A331R, A335R] mutant. The top panel shows the mScarlet-IFT54 signal, the middle panel displays the CFAP36-mStayGold signal, and the bottom panel presents a merged view of both signals. The strong central signal intensity is due to epi-fluorescence signal from the cell body. (G)Schematic model of the steps proposed in the retrieval of polyubiquitinated receptors from cilia by CFAP36•ARL3•GTP and retrograde IFT.

To further confirm that these interactions involve IFT70, we employed CRISPR/Cas9 to substitute two alanine residues in IFT70 (A331 and A335) with bulkier arginine residues, which are predicted to disrupt the interaction with the CFAP36 α8 helix (**Fig. 7B**). These mutations were introduced into the endogenous *Ift70* locus in the double-tagged IFT54-mScarlet;CFAP36-mStayGold strain (**Fig. S5C**) and validated by sequencing. TIRF microscopy analysis of this mutant strain revealed that mutating the predicted IFT70 interface did not interfere with IFT train movement, as judged by monitoring IFT54-mScarlet, but effectively decouples CFAP36 from retrograde transport (**Fig. 7F** and **Fig. S6G**). Collectively, our *in-vivo* imaging in strains with precisely targeted mutations support the coincidence detection model suggested by *in-silico* modeling.

## Discussion

In this study, we identified CFAP36 as a conserved, cilia-specific adaptor of intraflagellar transport (IFT) in both motile and primary cilia that functions to retrieve polyubiquitinated proteins. We discovered that CFAP36 can interact with a broad range of proteins in a ubiquitin-dependent manner, suggesting it plays a role as a general IFT adaptor of ubiquitinated proteins. Accordingly, we observed that depletion of CFAP36 disrupts retrieval of ubiquitinated transmembrane receptors and thereby Hh signaling in a murine cell line. Furthermore, we revealed the molecular mechanism by which it can simultaneously recognize polyubiquitin proteins and retrograde IFT trains. Collectively, our studies explain the functions of all the conserved structural features of CFAP36: the N-terminal BART domain binds ARL3•GTP to enhance recognition of polyubiquitin chains, helices α6 and α7 bind ubiquitin, helix α8 binds IFT70 paralogs of the IFT-B complex and the C-terminal helix α9 binds IFT144 of the IFT-A complex (**Fig. 7A**). By simultaneously binding subunits of the IFT-A and IFT-B complexes, CFAP36 uses a mechanism of coincidence detection that is only possible in retrograde trains because of a spatial arrangement distinct from anterograde trains to ensure exclusive coupling poly-ubiquitinated proteins to retrograde transport. A schematic of the proposed sequence of events is provided in **Fig. 7G**.

### Mechanism for recognizing polyubiquitinated substrates

Our proteomics experiments have demonstrated that CFAP36 is capable of binding a diverse clientele of polyubiquitinated proteins and ubiquitin linkage types. Its ability to recognize both K48- and K63-linked ubiquitin chains suggests that CFAP36 may retrieve all ubiquitinated protein from cilia and that the ubiquitin linkage type determines the subsequent fate of the cargo proteins, e.g. sorting into endosomes or proteasomal degradation.

The ability of CFAP36 to bind ubiquitin is dependent on helix α6 and the first UIM of helix α7; single-residue substitutions in either abolished ubiquitin binding *in vitro*. The intrinsically low affinity of CFAP36 for ubiquitin is substantially enhanced by ARL3•GTP. The ability of ARL3 to enhance ubiquitin binding in a nucleotide-dependent manner is not unique among the small GTPase family: the Golgi-localized ARF1•GTP enhances ubiquitin binding of GGA1 (Shiba *et al*., 2004) and RAB5, a regulator of early endosomal trafficking, enhances the ubiquitin-binding of its guanine nucleotide exchange factor, RABEX5, by releasing an autoinhibited state that obstructs its binding sites (Lauer *et al*., 2019).

ARL3•GTP may enhance the affinity of CFAP36 for ubiquitin chains through a similar mechanism of autoinhibition release or by altering the conformational dynamics of CFAP36 to favor ubiquitin binding. Alternatively, ARL3 could directly bind polyubiquitin chains, thereby enhancing the avidity of the interaction. All these scenarios would be consistent with our findings. We observed that C-terminally truncated CFAP36 exhibited enhanced binding to ubiquitin compared to full-length CFAP36 (**Fig. S3B**), with even greater binding observed upon the addition of ARL3•GTP. However, the precise mechanism by which ARL3•GTP enhances the recognition of polyubiquitin chains remains to be elucidated by further work and likely requires a structure of the CFAP36•ARL3•GTP•polyubiquitin complex.

ARL3 has an N-terminal amphipathic helix that promotes membrane binding in the GTP-bound state. In the context of cilia, the membrane localization of ARL3•GTP may bring it into proximity of ubiquitinated transmembrane proteins and therefore in a primed position to form a ternary complex with CFAP36. However, any membrane tethering by ARL3•GTP is likely to be transient, because the CFAP36 BART domain binds the hydrophobic face amphipathic helix (Lokaj *et al*., 2015) and is therefore likely to prevent or reduce membrane interaction of ARL3•GTP. Once ARL3•GTP is bound to CFAP36, the ternary complex would dissociate from the membrane, potentially facilitating interactions with polyubiquitin chains on transmembrane and soluble proteins, and the IFT machinery.

It remains unclear if ARL3•GTP travels with CFAP36 during retrograde IFT or if it is released from CFAP36 upon engaging with the IFT machinery. Live-cell TIRF microscopy of ARL3-YFP expressed in *C. reinhardtii* (Liu *et al*., 2022) found evidence only for movement of the protein by diffusion. However, the large pool of ARL3 present in flagella might have masked signal from the smaller pool of ARL3 undergoing retrograde transport. If ARL3 is indeed released at the stage of IFT coupling, the underlying mechanism remains to be determined.

The timing of ARL3 release has implications for the unloading of polyubiquitinated proteins from retrograde trains. If ARL3•GTP accompanies retrograde trains with CFAP36, an active mechanism for unloading might involve a GTPase-activating protein (GAP) at the ciliary base. A candidate GAP is RP2, a known ARL3 GAP (Veltel *et al*., 2008) that resides at the ciliary base (Evans *et al*., 2010) and is capable of disrupting pre-formed complexes of CFAP36•ARL3•GppNHp *in vitro*(Lokaj *et al*., 2015). RP2 or another GAP may displace CFAP36 from polyubiquitin chains through direct binding to ARL3•GTP, or by stimulating GTP hydrolysis and thereby generating an unstable complex (**Fig. 7G**). Alternatively, release could occur passively through the depolymerization of retrograde IFT trains into subcomplexes (i.e. IFT-A, IFT-B and dynein) at the ciliary base. Once freed from IFT trains and CFAP36, the polyubiquitinated proteins could then become substrates for deubiquitinases or other cellular pathways, including the ESCRT pathway (**Fig. 7G**). CFAP36 and ARL3 would then be free to diffuse.

### CFAP36 is necessary for Hedgehog signaling

Our study has identified CFAP36 as a novel and crucial component of the Hh signaling pathway. Depletion of CFAP36 disrupts the normal distribution of Hh pathway proteins in unstimulated IMCD3 cells and impairs their proper exit from cilia in response to stimuli. The resultant mislocalization ultimately prevents a transcriptional response, as evidenced by the failure to elevate *Ptch1* expression in CFAP36-depleted cells following SAG treatment. The observed hyperaccumulation of Smoothened, GPR161, and K63-linked ubiquitin chains in these cells serves as a clear indication that CFAP36 plays an integral role in the ubiquitin-dependent retrieval of signaling components from cilia.

The functionality of CFAP36 likely extends beyond Hh signaling. Given its critical role in retrieving polyubiquitinated proteins from cilia, it is plausible that CFAP36 is involved in other signaling pathways which rely on precise protein localization within cilia, such as phototransduction and Wnt and TGF-β signaling (Yildiz & Khanna, 2012; Christensen *et al*., 2017; Niehrs *et al*., 2025). In particular, CFAP36 might play a role in the weight-regulating leptin-melanocortin pathway based on recent findings demonstrating the ubiquitin-dependent removal of melanocortin receptor 4 (MC4R) and its interaction partner MRAP2 from cilia (Ojeda-Naharros *et al*., 2025). CFAP36 may also function in quality control pathways that maintain ciliary proteomes by removing mislocalized non-ciliary proteins following their ubiquitination. Consistent with a role beyond Hh signaling, CFAP36 is not a transcriptional target of the Hh signaling pathway (Daggubati *et al*., 2023) and the ciliary levels of the CFAP36 protein do not change following SAG-induced stimulation (May *et al*., 2021). Further investigations are needed to elucidate the full spectrum of cilia-dependent signaling and quality control pathways in which CFAP36 is involved.

Given that ciliopathies are often characterized by the dysregulation of ciliary signaling pathways and improper ciliary localization of proteins, CFAP36 is a compelling candidate for contributing to ciliopathies of previously unknown etiology. The potential involvement of CFAP36 in multiple cilia-dependent pathways suggests that mutations or dysregulations in this protein may affect various physiological processes. Mutations in its binding partner ARL3 are associated with retinal degeneration (Strom *et al*., 2016; Holtan *et al*., 2019) and Joubert syndrome (Alkanderi *et al*., 2018). Given these characteristics, CFAP36 merits further investigation to determine its genetic integrity in patients with ciliopathies.

### Coincidence detection allows specific recognition of retrograde trains

Our experimental data support a coincidence detection mechanism, in which CFAP36 needs to bind both IFT70 and IFT144 for robust transport, as mutation in either binding site abolished movement. The interaction is likely weak as live-cell imaging revealed CFAP36 occasionally detaches from trains but is subsequently reloaded onto later trains (train hopping), highlighting the presence of a fail-safe retrieval system. A similar process (carriage hopping) may also occur within an IFT train, which because of its polymeric nature contains multiple potential CFAP36 binding sites, but these dynamics are beyond the resolution of TIRF microscopy.

Our work also demonstrates the critical role of retrograde IFT train conformation in the retrieval of polyubiquitinated proteins from cilia. Defects that compromise the spatial organization of IFT-A and IFT-B subcomplexes within retrograde trains are likely to impair the coincidence detection mechanism used by CFAP36, thereby preventing the clearance of ubiquitinated proteins from cilia. Such disruptions may explain the Hh signaling defects following knockout of *Ift25* or *Ift27* in both mice and murine embryonic fibroblasts (Keady *et al*., 2011; Eguether *et al*., 2014). These mutations phenocopy knockdown of CFAP36, leading to the accumulation of Smo and Gpr161 in cilia and an attenuated transcriptional response to SAG (Keady *et al*., 2011; Eguether *et al*., 2014). The similarity of *Ift25/27* mutants to CFAP36 knockdown can be attributed to the role of the IFT25/27 heterodimer in retrograde transport where it mediates interactions between IFT-A and IFT-B (Lacey *et al*., 2024). Neither protein is resolved in a cryo-ET structure of the anterograde train (Lacey *et al*., 2023), which is consistent with their dispensability for anterograde transport and ciliogenesis (Keady *et al*., 2011; Eguether *et al*., 2014). Thus, our findings suggest that loss of IFT25 and IFT27 disrupt the coincidence detection mechanism vital for effective protein retrieval in cilia by interfering with proper retrograde train formation.

Missense mutations in CFAP36-binding sites could similarly disrupt coincidence detection. For example, the IFT70B A375V missense mutation associated with impaired Hh signaling in humans (Du *et al*., 2019), introduces a bulkier sidechain adjacent to the CFAP36 binding site. Molecular modeling predicts that this substitution distorts the spatial packing of the TPR repeats essential for CFAP36 binding, providing a potential structural basis for its pathogenicity.

### Mechanistic similarities with cytoplasmic dynein adaptors

CFAP36 shares similarities with the adaptor proteins of the cytoplasmic dynein•dynactin complex, which functions analogously to retrograde IFT trains to move cargos to the minus ends of microtubules. These cytoplasmic adaptors typically use α-helices to establish simultaneous interactions with both dynein and dynactin, while also possessing additional domains that bind small GTP-bound GTPases (Reck-Peterson *et al*., 2018). This is similar to how CFAP36 binds IFT-A, IFT-B and ARL3•GTP. The identification that ARL3•GTP enhances the ubiquitin-binding activity of CFAP36 further strengthens the role of GTPases as important regulators of both intracellular and intraflagellar transport.

Many adaptors modulate the motile properties of dynein, either enhancing or inhibiting movement. In our observations, IFT trains carrying CFAP36 travel at slower velocities than trains without. While CFAP36 might directly suppress retrograde transport, the reduced speed may also be the result of the additional drag of pulling transmembrane proteins through the ciliary membrane.

Despite these structural and functional similarities, a critical distinction exists between how ubiquitinated cargos are handled in intracellular and intraflagellar transport. In cilia, ubiquitinated proteins appear to be recognized directly by an adaptor and linked to the dynein-driven transport machinery. In contrast, in the cytoplasm, ubiquitinated proteins are first sequestered into endosomes, which are then recognized by adaptors for dynein-mediated transport. Nevertheless, the structural parallels between CFAP36 and dynein•dynactin adaptors suggest that comparable, non-endosomal mechanisms of actively transporting ubiquitinated proteins may be present in cytoplasmic transport.

### Relationship with the proposed TOM1L2-BBSome pathway

Notably, the CFAP36-mediated mechanism for retrieving polyubiquitinated proteins from cilia identified here is distinct from the previously proposed TOM1L2-mediated mechanism (Shinde *et al*., 2023), as it requires neither TOM1L2 nor the BBSome. This is surprising as there is ample evidence linking TOM1L2 and the BBSome to the export of ciliary proteins from cilia. For example, loss of BBSome subunits and its GTPase partner, ARL6, cause the accumulation of ciliary ubiquitin and ciliary levels of proteins that would normally have low basal levels. These include Phospholipase D in *C. reinhardtii* (Lechtreck *et al*., 2013, 2009), dopamine receptor 1 in mice (Domire *et al*., 2011; Zhang *et al*., 2013), and Smoothened and GPR161 in mammalian cell lines (Seo *et al*., 2011; Zhang *et al*., 2011; Desai *et al*., 2020). Evidence for the involvement of TOM1L2 comes from the ciliary accumulation of K63-linked ubiquitin and ciliary proteins normally removed from cilia (e.g. Smoothened, GPR161 and SSTR3) following genetic disruption in cell lines (Shinde *et al*., 2023). Biochemical interactions between recombinant TOM1L2 and the BBSome led to the model where TOM1L2 and the BBSome function together as the adaptor linking ubiquitinated proteins to the IFT machinery (Shinde *et al*., 2023).

Despite the similar phenotypes following CFAP36 and TOM1L2 knockdown in cell lines, the TOM1L2 pathway is unlikely to be fully redundant with the CFAP36 pathway. If it were, the knockdown of either factor would not result in the mislocalization of K63-linked ubiquitin or Hh-pathway-related proteins. We therefore propose that TOM1L2 functions downstream of CFAP36 at the ciliary base, potentially as a mechanism for linking polyubiquitinated cargos to the ESCRT pathway. Further work is needed to test this hypothesis.

## Summary

In summary, we have identified and validated a novel, highly conserved mechanism for the recognition and retrieval of polyubiquitinated protein from cilia and flagella. The identification of CFAP36•ARL3 defines a ubiquitin-dependent IFT adaptor and identifies a new role for ARL3 as a cargo loading factor for retrograde transport. The identified coincidence detection mechanism demonstrates how IFT train reconfiguration at the ciliary tip enables selective recognition of retrograde IFT trains. These findings enhance our understanding of organelle-specific mechanisms of ubiquitin-mediated protein transport and homeostasis.

### Limitations

We have demonstrated that ARL3•GTP enhances the recognition of polyubiquitin chains by CFAP36 but not the mechanism by which this is achieved. We do not yet understand the timing of ARL3•GTP release: is it offloaded immediately following cargo-binding or does it traffic with CFAP36 on retrograde IFT trains? If it does migrate with CFAP36, is GTP hydrolysis and nucleotide exchange part of the mechanism that releases polyubiquitinated cargos from IFT trains? Does this release occur before or after the transition zone, and what is the fate of the polyubiquitinated proteins following uncoupling from CFAP36 and IFT trains? Does TOM1L2 and/or the BBSome function downstream of CFAP36-mediated retrieval? Addressing these questions will enhance our mechanistic understanding of this pathway.

## Acknowledgements

IMCD3-FlpIn cells were a gift from Peter Jackson (Stanford University). Ubiquitin ligation enzymes were a gift from the MRC-PPU Reagents and Services. Fluorescence microscopy data were collected at the Harvard Medical School (HMS) Core for Imaging technology and Education (CITE). SEC-MALS data were collected at the HMS Center for Macromolecular Interactions (CMI). Portions of this research were conducted on the O2 High Performance Compute Cluster, supported by the Research Computing Group at HMS. We thank E.W. Schmid for advice on evaluating AlphaFold2-Multimer predictions using SPOC, and M. Bao, R. Subramanian and Y. Kulathu for comments on the manuscript.

## Funding

SML was supported by a Sara Elizabeth O’Brien Trust Postdoctoral Fellowship awarded through the Charles A. King Trust Postdoctoral Research Fellowship Program. AB was supported by the Smith Family Foundation and NIGMS grants R01GM141109 and R01GM143183.

## Author contributions

SML performed all experiments. RJE performed quantitative mass spectrometry. AB supervised the research. SML and AB wrote the manuscript.

## Competing interests statement

The authors declare no competing interests.

## Materials and Methods

### Materials availability

Wild-type *Chlamydomonas reinhardtii* (CC-1690), mScarlet-IFT54 (CC-5860), and strains generated in this study (mScarlet-IFT54 CFAP36-mStayGold (CC-6258), mScarlet-IFT54 CFAP36[L438E, K442E]-mStayGold (CC-6259), mScarlet-IFT54 CFAP36[A493E, K495A]-mStayGold (CC-6260), mScarlet-IFT54 CFAP36[L438E, K442E, A493E, K495A]-mStayGold (CC-6261), mScarlet-IFT54 CFAP36-mStayGold IFT70[A331R, A335R] (CC-6262)) are available from the Chlamydomonas Resource Center (University of Minnesota). Plasmids generated during this work have been deposited to Addgene (https://www.addgene.org) with IDs 238214 to 238223.

### Data and code availability

- Quantitative mass spectrometry data has been deposited to PRIDE with accession code PXD062789.
- Any additional information required to reanalyze the data reported in this paper is available from the lead contact upon request.

### Experimental models and organisms

Wild-type and mScarlet-IFT54 *C. reinhardtii* strains (CC-1690, CC-5860) were obtained from the Chlamydomonas Resource Center. All strains were cultured in TAP media (20 mM Tris base, 0.62 mM K_2_HPO_4_, 0.41 mM KH_2_PO_4_, 7.5 mM NH_4_Cl, 0.4 mM MgSO_4_, 0.34 mM CaCl_2_, 0.1% v/v glacial acetic acid, 150 µM EDTA, 77 µM ZnSO_4_, 184 µM H_3_BO_3_, 25.6 µM MnCl_2_, 6.8 µM CoCl_2_, 9.8 µM CuSO_4_, 0.9 µM (NH4)_6_Mo_7_O_24_, 18 µM FeSO_4_, adjusted to pH 7.0 with HCl). Cells were cultured in 14:10 h light:dark cycles.

*Leishmania tarentolae* strain P10 (Jena Bioscience, #LT-101) cells were grown in BHI medium (HIMEDIA, #N210) containing 5 μg/mL hemin chloride (Sigma, #3741) and 10 U/mL penicillin-streptomycin (Gibco, #15070063) at 26 °C in the dark. Cells were maintained as static suspension cultures as described in the LEXSY expression kit manual (Jena Bioscience).

### Cell lines

IMCD3-FlpIn cells were a gift from Peter Jackson (Stanford University) (Mukhopadhyay *et al*., 2010). IMCD3 cells were cultured in DMEM:F12 (Gibco, #11320033) supplemented with 10% FBS (Sigma, #12306C), 2 mM L-glutamine (Gibco, #25030081), and 100 U/ml penicillin/streptomycin (Gibco, #15070063) subsequently referred to as complete DMEM:F12. All cells were maintained at 37 °C in a humidified incubator with 5% CO_2_ and routinely passaged every 2-3 days.

## Methods

### Construction of expression plasmids

Gene fragments were synthesized (TwistBio) or amplified by polymerase chain reaction (PCR) from *C. reinhardtii* cells processed with QuickExtract DNA Extraction Solution (Biosearch Technologies, #QE0905T) using Platinum SuperFi II polymerase (Thermo Fisher Scientific, #12368010) in the presence of 1 M betaine (Sigma, #B0300). Gene fragments were inserted into pET15 expression plasmids using Gibson Assembly cloning (NEB, #E2621).

### Site-directed mutagenesis

Point mutations were introduced by PCR using KOD One polymerase (Toyobo, #KMM-201) and primers designed with the QuikChange Primer Design Program (Agilent).

### Protein expression and purification

Recombinant proteins were expressed in *E. coli* host strain Rosetta 2 (Sigma, #71402) grown in 2xYT medium (RPI, #X15600) supplemented with antibiotics (100 µg/mL carbenicillin or 50 µg/mL kanamycin, as appropriate). Expression cultures with added autoinduction mix (0.6% v/v glycerol, 0.05% w/v glucose, 0.2% w/v lactose) were inoculated with 2% v/v freshly saturated 2xYT overnight culture and incubated at 18–25 °C for 24 h at 180 rpm shaking speed. Cells were harvested by centrifugation at 4,200 × *g* and 4 °C for 15 min and resuspended in 40 mL lysis buffer per liter expression culture (50 mM Tris pH 7.5, 300 mM NaCl, 0.5 mM TCEP, 1 mM 4-(2-aminoethyl)benzene-1-sulfonyl fluoride (AEBSF; VWR, #102988-758), 1 mM benzamidine, 10 µM leupeptin (VWR, #102988-516)). Bacterial cells were lysed with three passes on a Microfluidizer LM10 at 17,000 psi and subsequently clarified by centrifugation at 35,000 × *g* and 4 °C for 30 min.

His-tagged proteins were purified on nickel-nitrilotriacetic acid (Ni-NTA) agarose (Thermo Scientific, #88222) equilibrated with His-wash buffer (50 mM Tris pH 7.5, 300 mM NaCl, 0.5 mM TCEP, 10 mM Imidazole pH 8.0) in a gravity flow column. The resin was extensively washed and bound proteins eluted with His-elution buffer (50 mM Tris pH 7.5, 300 mM NaCl, 0.5 mM TCEP, 250 mM Imidazole pH 8.0). Eluted proteins were mixed with Human Rhinovirus (HRV) 3C Protease and dialyzed overnight in 50 mM Tris pH 8.5, 50 mM NaCl at 4 °C overnight, and subsequently reapplied to equilibrated Ni-NTA resin. The protein in the flow-through was applied to an ion-exchange chromatography MonoQ 5/50 column (Cytiva) equilibrated with 50 mM Tris pH 8.5, 50 mM NaCl and eluted using a NaCl salt gradient. As a final purification step, proteins were applied to size exclusion chromatography on a Superdex 200 Increase 10/300 column (Cytiva) equilibrated in 50 mM Tris pH 7.5, 150 mM NaCl, 0.5 mM TCEP.

Bacterial cell pellets of ubiquitin and AVI-tagged ubiquitin expression were resuspended in 20 mL Ub lysis buffer (1 mM EDTA, 1 mM AEBSF and 1 mM benzamidine) per liter of expression culture and lysed by sonication. Subsequently, 100 mM sodium acetate pH 4.5 was added and the lysate incubated for 3 h at 20 °C. The lysate was diluted to 50 mM sodium acetate through the addition of water and then clarified by centrifugation at 35,000 × *g* and 4 °C for 30 min. Ubiquitin was purified by ion-exchange chromatography on Source 15S resin (Cytiva, #17-0944-10) equilibrated with 50 mM sodium acetate pH 4.5 and eluted by NaCl salt gradient. Eluted ubiquitin fractions were supplemented with 100 mM Tris-HCl pH 8.5, concentrated in centrifugal filter units (Amicon) and finally buffer-exchanged into 50 mM Tris-HCl pH 7.5.

### Ubiquitin chain ligation and biotinylation

Ubiquitin chains were assembled from 1.5 mM ubiquitin in 40 mM Tris-HCl pH 7.5, 10 mM MgCl_2_ and 10 mM ATP at 30 °C for 2-3 h. AVI-tagged ubiquitin chains were assembled from a mixture of 1.5 mM ubiquitin and 0.3 mM ubiquitin[1-74]-AVI at 30 °C for 16 h. K48-linkages were assembled with 1 µM UBE1 and 25 µM UBE2R1, and K63-linkages with 1 µM UBE1, 20 µM UBE2N and 20 µM UBE2V1. Ubiquitin chains were separated by length using ion exchange chromatography on Source 15S media (Cytiva #17-0944-10) in 50 mM Sodium acetate pH 4.5 using a NaCl gradient. Th pH of the eluted fractions was adjusted with 100 mM Tris-HCl pH 8.5 prior to concentration in centrifugal filter units (Amicon) and finally buffer exchanged into 50 mM Tris-HCl pH 7.5. Biotinylation of 200 μM Avi-tagged Ub chains was performed with 1 μM BirA in 50 mM Tris-HCl pH 7.5, 5 mM MgCl_2_, 2 mM ATP and 600 μM biotin for 2 h at 25 °C, and subsequent buffer exchange into 50 mM Tris-HCl pH 7.5 to remove free biotin.

### Size Exclusion Chromatography with Multi-Angle Light Scattering (SEC-MALS)

Recombinant full-length porcine CFAP36 was subjected to SEC-MALS on a Agilent 1260 Infinity LC System equipped with Wyatt Dawn Heleos Multi-Angle Light Scattering (MALS, 18-angle) detector, with in-line DLS detector and Optilab TrEX refractive index detector. Approximately 50 µL protein at 1 mg/mL was loaded on a SEPAX SRT SEC-150 column equilibrated with degassed HBS buffer (25 mM HEPES pH 7.4, 150 mM NaCl, 0.25 mM TCEP) at a 0.5 ml/min flow rate. Bovine Serum Albumin (BSA) was used as a calibration standard.

### Flagella/cilia isolation

All flagella isolation procedures were conducted at 4 °C. The method for flagella isolation from *C. reinhardtii* was modified from the dibucaine method from Craige et al. (Craige *et al*., 2013). Briefly, CC-1690 cells were grown in 8 L TAP, harvested and washed twice with 10 mM HEPES pH 7.4. Cells were resuspended in ice-cold HMS buffer (10 mM HEPES, 5 mM MgSO_4_, 4% sucrose). Their flagella were detached using 4.2 mM dibucaine and pipetting cells up and down 10 times using a 10-mL serological pipette, followed by addition of 0.5 mM EGTA to quench the reaction after 2 min. Deflagellated cell bodies were removed by centrifugation at 1200 × *g* for 3 min. Remaining cell bodies were separated by centrifugation on a 25% sucrose cushion at 1700 × *g* for 10 min. Flagella were then sedimented by centrifugation at 12,000 × *g* for 15 min and stored at -80 °C until further use.

The method for flagella isolation from *L. tarentolae* was adapted from Beneke et al. (Beneke *et al*., 2020). Briefly, *L. tarentolae* strain P10 (Jena Bioscience, #LT-101) were grown in BHI medium (HIMEDIA, #N210) supplemented with 5 mg/mL hemin chloride (Sigma, #3741) and 10 U/mL Penicillin-Streptomycin (Gibco, #15070063) at 26 °C. A static suspension culture in late logarithmic growth phase was used in 1:10 dilution to inoculate a 300 mL suspension culture shaking at 120 rpm and grown overnight. The next day, cells were collected by centrifugation at 800 × *g* for 15 min, washed once with PBS and resuspended in 20 mL Lt-PIPES buffer (10 mM PIPES pH 7.5, 10 mM NaCl, 75 mM CaCl_2_, 1 mM MgCl_2_, 0.32 M sucrose, 1x protease inhibitor (Sigma, #S8820)). Flagella were detached by mechanical shearing with 100 passes through a 22-gauge needle attached to a 10 mL syringe and then separated from cell bodies by sucrose gradient centrifugation (layers of 33, 53, and 63% sucrose) at 800 × *g* for 15 min. Flagella were further purified by centrifugation on a 33% sucrose cushion at 100,000 × *g* for 1 h and stored at -80 °C until further use.

The purification of cilia from porcine trachea (*Sus scrofa*) was adapted from the method for bovine respiratory cilia isolation described in Gui et al. (Gui *et al*., 2021). A total of eight fresh porcine trachea (Animal Technologies, Inc.) were shipped on wet ice overnight, trimmed from excess tissue and washed three times each for 5 min with 20 mL ice-cold PBS under gentle agitation. One end of the trachea was closed off with plastic wrap and the trachea filled with 15 mL cilia-extraction buffer (20 mM PIPES pH 5.5, 20 mM CaCl_2_, 250 mM sucrose, 1x protease inhibitor, 0.01% Triton-X100). Cilia were detached by 10 gentle strokes with an inserted soft nylon brush. The suspension was collected and combined with a second wash-out of the trachea (2 min gentle agitation) and wash-out from the brush. The combined sample was passed through a 50 µm mesh tissue strainer (pluriSelect, #43-50050-01) to remove larger tissue pieces and then centrifuged at 1000 × *g* for 5 min to remove smaller cell debris. The supernatant was placed on a 60% sucrose cushion and centrifuged at 4200 × *g* for 15 min. The cilia-enriched fraction was collected from the cushion surface, diluted 1:10 in cilia-extraction buffer without sucrose, pelleted by centrifugation at 40,000 × *g* for 30 min and stored at -80 °C until further use.

### Pulldown with biotinylated ubiquitin from ciliary lysates

Frozen isolated flagella (from *L. tarentolae* or *C. reinhardtii*) were briefly thawed in a 20 °C water bath, transferred to ice and resuspended in 0.5 mL ice-cold HDMS supplemented with 1% NP-40 (Thermo, cat# 28324), then lysed by end-over-end agitation for 30 min at 4 °C. Lysates were clarified by centrifugation at 17,000 × *g*, 4 °C for 10 min and supernatants diluted with 0.5 mL ice-cold HDMS without NP-40. Per pulldown, 50 µL of Streptavidin agarose beads (Thermo Fisher, #20353) were centrifuged at 500 × *g* for 1 min to remove storage solution and washed twice with 20 volumes of wash buffer PW (10 mM Tris-HCl pH 7.4, 150 mM NaCl, 1x protease inhibitor cocktail (Sigma, #S8830)). The beads were pelleted and resuspended in 200 µL wash buffer containing 100 µg biotinylated ubiquitin chains (per 50 µL beads) at 20 °C for 30 min rotating end-over-end at approximately 30 rpm. The beads were washed three times with 1 mL wash buffer and once with 1 mL wash buffer supplemented with 10 µM Biotin to block free binding sites. The beads were resuspended as a 50% slurry in wash buffer, aliquoted as 100 µl into low-binding 1.5 mL tubes and incubated with the clarified flagellar lysate rotating end-over-end at 4 °C for 2 h. The beads were then washed a total of four times with 1 mL wash buffer and transferred to a clean 1.5 mL low-binding tube for the final wash. The beads were resuspended in 200 µL elution buffer (10 mM Tris-HCl pH 7.5, 10% SDS) and heated at 70 °C for 10 min. After centrifugation at 1,000 × *g* for 2 min, the supernatant was transferred to a clean low-binding 1.5 mL tube and 15 µL of each sample was analyzed by SDS-PAGE and silver stain.

### Pulldown with ARL3-CFAP36 fusion

Recombinant fusion protein of porcine ARL3-4x(GS)-3xFlag-4x(GS)-CFAP36_2-342_-10His with either wild-type CFAP36 or ubiquitin-binding mutant S163E were immobilized on anti-Flag resin equilibrated in GTP-wash buffer (10 mM Tris pH 7.4, 150 mM NaCl, 1x protease inhibitors, 0.02% NP-40, 100 µM GTP, 200 µM MgCl_2_) using 60 µg of bait protein and 50 µl of resin per pulldown for 30 min at 22 °C with gentle end-over-end rotation. Lysates of frozen isolated porcine cilia were prepared as described above for *L. tarentolae* or *C. reinhardtii* flagella. The resin was washed twice with 1 mL GTP-wash buffer and subsequently incubated with the clarified ciliary lysate for 2 h at 4 °C with gentle end-over-end rotation. Each pulldown was washed four times with 1 mL GTP-wash buffer and the beads transferred to a clean 1.5 mL tube in the final wash step. Next, the resin was pelleted at 500 × *g* for 2 min and resuspended in 200 µL elution buffer (10 mM Tris pH 7.4, 10% SDS) and incubated at 70 °C for 10 min. The samples were centrifuged at 1000 × *g* for 2 min and the supernatant collected in a clean 1.5 mL tube.

### Sample preparation for quantitative mass spectrometry with tandem mass tagging (TMT-MS)

Equal amounts of hydrophobic and hydrophilic magnetic SP3 beads (Cytiva, #44152105050250 and #24152105050250) were resuspended in HPLC-grade water (Thermo, #047146) at 50 mg/mL and 0.2 mg of SP3 beads were added to each pulldown supernatant. Binding was induced by addition of 1 volume ethanol and the bead suspensions were incubated at 24 °C for 5 min shaking at 1000 rpm in a Thermomixer (Eppendorf). The beads were washed a total of three times with 1 mL 80% ethanol in HPLC-grade water and transferred to a clean 1.5 mL low-binding tube for the final wash. The beads were stored dry at - 80 °C for further processing.

On bead digestion was performed by adding 50 µL 200 mM 3-[4-(2-Hydroxyethyl)piperazin-1-yl]propane-1-sulfonic acid (EPPS), pH8.5 with 4% acetonitrile with 1 µL of 2 mg/mL Lys-C (Wako, #129-02541) and shaking at 1000 rpm at 37 °C for 3 hours. Another 50 µL 200 mM EPPS was added with 1 µL trypsin (Promega, #V5113) and digestion continued overnight at 37 °C with shaking at 1000 rpm then the digests were transferred to new tubes. To each digest, 30 µL of liquid chromatography–MS (LC-MS) grade acetonitrile was added and the samples were labeled with either TMTpro or TMT11plex (Thermo Fischer Scientific, #A52045 or #A34807). Once labeling was determined to be greater than 95% by LC-SPS-MS3, samples were quenched by adding hydroxylamine to a final concentration of 0.5% v/v for 15 min at room temperature. The reactions were acidified with formic acid and pooled. The multiplexes were fractionated by high pH reversed-phase benchtop fractionation (Thermo Scientific, #84868) using the following step gradient of 10%, 12.5%, 15%, 17.5%, 20%, 25%, 30%, 35%, 40%, 50%, 65%, and 80% acetonitrile. Fractions 1 and 7, 2 and 8, 3 and 9, 4 and 10, 5 and 11, and 6 and 12 were pooled to generate 6 final fractions, which were dried, and desalted by StageTip.

### TMT-MS data collection, analysis and visualization

Data were collected using an Orbitrap Eclipse mass spectrometer (Thermo Fisher Scientific) paired with a Vanquish Neo high-performance liquid chromatography system (Thermo Fisher Scientific) with or without field asymmetric ion mobility spectrometry (FAIMS). Peptides were separated on an EASY-Spray PepMap column (Thermo Scientific, #ES75750PN) using either a three or four hour gradient of water to acetonitrile with 0.1% formic acid then analyzed with a synchronous-precursor-selection (SPS)–MS_3_ acquisition method. The high-resolution MS1 spectra were first collected in the orbitrap (resolution 120,000; a scan range of 400-1500; automated gain control (AGC) target of 200%, and dynamic exclusion 60 sec). Peptides were fragmented using collision-induced dissociation (CID, CE=35%) and MS2 scans were collected in the ion trap and AGC and maximum injection time set to standard and auto, respectively. Real-time search and FAIMS were used with static modifications of alkylated cysteine (+57.0215) and TMTpro at the N-terminus and lysine sidechains (+304.2071) and dynamic modification of oxidized methionine (+15.9949) were permitted. Precursors were fragmented using high-energy collision-induced dissociation (HCD, CE=50% and 60% for TMTpro and TMT11plex, respectively) then analyzed in the orbitrap with a resolution of 50,000.

The raw spectra were converted to mzXMLand subsequently post-search calibrated. Peptide spectrum matches (PSMs) were made using COMET-based searches (v2019.01 rev. 5)(Eng *et al*., 2013) with TMT set as a static modification (TMTpro and TMT11 set as +304.2071 and +229.162932, respectively). Alkylated cysteine was also set as a static modification (+57.02146) and oxidated methionine was set as a dynamic modification (+15.9949). The protein databases also contained common contaminants and reversed sequences as decoys. The mass tolerance was set to 20 ppm and fragment ion tolerance was set to 1.00. Using a target-decoy strategy, PSMs were filtered by linear discriminant analysis (LDA) with a false discovery rate (FDR) of 1% and an FDR of 1% for collapsed proteins. For TMT quantification data were filtered for an isolation specificity above 70% and a minimum summed signal-to-noise of 200 across all TMT channels.

Quantified TMT-MS data were normalized using total peptide amount and scaled to the average of all samples. For heatmaps, the scaled data was log2-transformed and a one-way ANOVA performed for each protein across experimental groups, followed by Benjamini-Hochberg FDR correction for multiple testing. Proteins with q-values below 0.01 were considered statistically significant, and those exhibiting a fold-change greater than 2 compared to the control group were retained. To cluster the data for visualization, a Principal Component Analysis (PCA) was performed and proteins sorted based on the first principal component values. For volcano plots, the unscaled intensities were log2(x+1) transformed and comparison of experimental groups was conducted using pairwise Student’s t-tests. The resulting p-values were adjusted for multiple testing using Benjamini-Hochberg FDR correction and log2 fold changes were calculated as the difference between mean group values of the transformed data. Human homologs were identified with phmmer (v3.3.2) and BLASTp (blast.ncbi.nlm.nih.gov). For the ubiquitin pulldowns, known contaminants (including ACACA, ECI1, PCCA, and PCCB), bait proteins (UBC), and proteasomal subunits (starting with PSMD, PSMC, and PSMA) were removed from the dataset. Data analysis was performed using the scipy (v1.13.1) python library. The mass spectrometry proteomics data have been deposited to the ProteomeXchange Consortium via the PRIDE (Perez-Riverol *et al*., 2024) partner repository with the dataset identifier PXD062789.

### Ubiquitin Interacting Motif (UIM) sequence motif

Putative UIM motifs in human CFAP36 were identified using remote homology detection with HHpred (Zimmermann *et al*., 2018). Consensus sequence motifs of UIMs were generated from annotated motifs of PROSITE (Sigrist *et al*., 2013) database entry PS50330 using WebLogo3 (Crooks *et al*., 2004).

### ScanNet Analysis

Protein-protein interface probabilities of human CFAP36 were generated from the AlphaFold Database (Varadi *et al*., 2023) entry AF-Q96G28-F1-v4 using ScanNet-PPI (Tubiana *et al*., 2022). Protein structure alignments and visualizations were produced in ChimeraX (v1.8) (Meng *et al*., 2023).

### AlphaFold2-Multimer screen

For the AlphaFold2-Multimer screen, gene names of ciliary proteins were retrieved from the CilioGenics database v0.2.3 (mean score ≥1.0) (Pir *et al*., 2024), SYSCILIA gold standard (SCGSv2) (Vasquez *et al*., 2021) and the UniProt database (based on cilia- and sperm-related annotations and GO terms) (Consortium *et al*., 2022). In addition, we added ciliary genes identified by structural proteomics in the Brown lab, and for the CFAP36 screen also ubiquitin and top entries from COXPRESdb (v8) (Obayashi *et al*., 2022) and STRINGdb (Szklarczyk *et al*., 2022) for both TOM1L2 and CFAP36. Genes were mapped to UniProt IDs and merged yielding a final list of 3379 unique entries. The lists for screens against IFT70B and IFT144 excluded the COXPRESdb and STRINGdb entries and totaled 2780 unique entries. Protein sequences were retrieved from UniProt and pair-wise structure predictions were conducted using multiple sequence alignments generated by the MMseqs2 server and AlphaFold2-Multimer (v2.3.2) (Jumper *et al*., 2021; Evans *et al*., 2021) via localcolabfold (v1.5.5) (Mirdita *et al*., 2022) on the O2 High Performance Compute Cluster at Harvard Medical School with three recycles, five models and no templates. Compact Structure Prediction and Omics informed Classifier (cSPOC) scores (Schmid & Walter, 2025) were calculated from the top three ranked predictions per complex. The predictions were sorted by descending cSPOC scores. The elbow point in the distribution of scores was detected using the KneeLocator algorithm (Satopää *et al*., 2011) of the kneed (v0.8.5) python library.

### Small interfering RNA (siRNA) knockdown, ciliation and Smoothened agonist (SAG) treatment

Four acid-washed 12 mm glass coverslips stored in 100% ethanol were placed per well of a 6-well plate and allowed to dry. Next, 0.2 Mio IMCD3 cells in complete DMEM:F12 were seeded per well and incubated overnight at 37 °C. Per transfection, 10 µL DharmaFECT1 transfection reagent (Dharmacon, #T-2005-01) was mixed with 190 µL Opti-Mem at 23 °C and incubated for 5 min at 23 °C. Meanwhile, 7.5 µl of 20 µM siRNA, either against CFAP36 or non-targeting control (Dharmacon, #L-049797-01 and #D-001810-01), was mixed with 192.5 µL Opti-MEM. Transfection reagent and siRNA mixtures were combined and incubated for 15 min at 23 °C. The cells were washed once with PBS, the medium was exchanged to complete DMEM:F12 without Pen/Strep and the transfection mix added dropwise. The next day, the cells were washed once with PBS, the medium exchanged to complete DMEM:F12 without FBS to induce ciliation and the cells incubated for 24 h. Cells were treated with 200 nM SAG (Sigma, Cat #SML1314-1MG) for 2 h prior to fixation.

### Fixation, immunofluorescent staining and imaging

IMCD3 cells were washed briefly with 2 mL PBS and then fixed with 2 mL 4% Formaldehyde and 4% sucrose in PBS for 10 min at 23 °C. Coverslips were washed once with 2 mL PBS + 0.2% Triton X-100 (Sigma, #T8787) for 5 min and three times with 2 mL PBS for 3 min each at 23 °C, then incubated for 1 h in blocking buffer (4% normal donkey serum in PBS) followed by 2 h incubation with primary antibodies diluted in PBS + 1% bovine serum albumin (BSA) at 23 °C. Cells were washed three times with 2 mL PBS for 5 min each, incubated for 1 h with secondary antibodies diluted in PBS + 1% BSA at 23 °C and washed again three times with 2 mL PBS for 3 min each. Coverslips were carefully blotted from the side and mounted on microscope slides with 10 µl Fluoromount G (SouthernBiotech, #0100-01) and sealed with clear nail polish. Slides were imaged on a Nikon Ti motorized inverted microscope equipped with Yokagawa CSU-X1 spinning disk confocal, a Nikon LUN-F XL solid state laser combiner (405, 488, 561, 640 nm) and Hamamatsu ORCA-Fusion BT sCMOS camera. Primary antibodies: Mouse anti-ARL13B (Proteintech, #50-173-7149, 1:500), Rabbit anti-SMO (Proteintech, #20787-1-AP, 1:100), Rabbit anti-GPR161 (Proteintech, #13398-1-AP, 1:100), Rabbit anti-Ub K63-secific (Sigma, #05-1308, 1:200). Secondary antibodies: Goat anti-mouse Alexa Fluor 488 (Invitrogen, #A32723, 1:500), donkey anti-rabbit Alexa Fluor 568 (Abcam, #ab175470, 1:500), goat anti-rabbit Alexa Fluor 680 (Invitrogen, #A21109, 1:500). Visualizations of representative cilia were prepared using OMERO webclient (https://www.openmicroscopy.org/) (Allan et al., 2012) using consistent intensity range settings across treatment conditions.

Z-stacks of ciliated cells were analyzed using the CiliaQ plugin pipeline (https://github.com/hansenjn/CiliaQ) (Hansen et al., 2021) for ImageJ/Fiji (Schindelin et al., 2012). Cilia were automatically segmented using the ciliary marker channel (ARL13B) with CiliaQ preparator (v0.1.2) and Canny 3D Thresholder (Gauss sigma 1.0, Canny alpha 5.0, Low/High threshold method: Triangle/Otsu). Identified objects were filtered by minimum voxel size of 1750 to remove noise and manually inspected with CiliaQ Editor (v0.0.3) to remove any remaining incorrectly identified objects. Subsequently, signal intensities of target proteins within cilia and dimensions of individual cilia reconstructions were analyzed using CiliaQ (v0.1.7) (0.065 µm/px, voxel depth 0.2 µm).

### Quantitative reverse transcription polymerase chain reaction (RT-qPCR)

IMCD3 cells in 6-well plates were transfected with siRNA and treated with SAG as described above. Cells were briefly washed with PBS, dissociated using trypsin and harvested in 1 mL PBS. Cells were collected by centrifugation at 300 × *g* for 5 min and washed once with 1 mL PBS. Total RNA was isolated using SV Total RNA Isolation System (Promega, #Z3101) following the manufacturer’s instructions, flash frozen in liquid nitrogen and stored at -80 °C until further use. cDNA was synthesized using LunaScript RT SuperMix Kit (NEB, #E3010) according to the manufacturer’s instructions using random hexamers. RT-qPCR was performed with a SYBR Green master mix (1.7% v/v glycerol, 12.76 mM Tris-HCl pH 8.0, 53.2 mM KCl, 5.32 mM MgCl2, 0.21% v/v Tween 20, 212.7 mg/mL BSA, 0.71 × SYBR Green (Sigma, #T8531) and run on a Bio-Rad CFX384. TATA-box-binding protein (TBP) and Large ribosomal subunit protein eL27 (RPL27) served as reference genes. Patched 1 mRNA levels were calculated using ΔΔCq method relative to RPL27. Differences were assessed using Tukey’s HSD test with a significance level of α = 0.05.

### Western Blot

Cell lysates were quantified with a BSA standard curve using Bradford dye (BioRad, #5000205) by absorbance at 595 nm and run on SDS-PAGE at 180 V for 30 min. Proteins were then wet-transferred from SDS-PAGE gels in Tris-Glycine transfer buffer (383 mM glycine, 50 mM Tris base, 10% ethanol) to nitrocellulose membranes (Cytiva, #10600002) at 90 V for 90 min in a cooled transfer tank. Membranes were blocked with 5% BSA in TBS (32 mM Tris base, 150 mM NaCl) for 1 h at 23 °C and then incubated with primary antibodies diluted in 5% BSA in TBS for 2 h at 23 °C or overnight at 4 °C. Membranes were washed 3 times with TBS-T (TBS + 0.1% Tween-20) for 15 min each and then incubated with secondary antibodies diluted in 5% BSA in TBS for 1 h, then washed 3 times as above. Fluorescently labeled secondary antibodies were imaged on Licor Odyssey CLx in 700 nm and 800 nm channels and analyzed in ImageStudioLite (v5.2.5). Primary antibodies: rabbit anti-CFAP36 (Proteintech, #24276-1-AP, 1:5000), mouse anti-acetylated α-tubulin (Invitrogen, #32-2700, 1:2500). Secondary antibodies: goat anti-rabbit Alexa Fluor 680 (Invitrogen, #A21109, 1:10,000), goat anti-mouse Alexa Fluor 800 (Invitrogen, #A32730, 1:10,000).

### CRISPR/Cas9-mediated gene editing

The experimental design for the knock-in of mStayGold and point mutations into endogenous *C. reinhardtii Cfap36* and *Ift70* loci is detailed in **Fig. S5** and derived from the method developed by Nievergelt and coworkers (Nievergelt *et al*., 2023). Technical difficulties with modifying the *Ift144* locus prevented interface-interfering mutations being introduced into IFT144. CRISPR/Cas9 target sites on C. reinhardtii CC4532 (v6.1) genome (Craig et al., 2022) were selected using the CHOPCHOP web tool (https://chopchop.cbu.uib.no/) (Labun et al., 2019). Donor template plasmids (5 µg) were digested with BspQI (NEB, #R0712S) for 1 h at 50 °C and purified in spin columns and eluted at ∼400 ng/µL final concentration (Zymo Research, #D4033). Synthetic single guide RNA (SafeEdit sgRNA, GenScript) were resuspended to 100 µM and 2 µL mixed with 1.8 µL Cas9 and 16 µL duplex buffer (IDT, #1081058) at 20 °C to form ribonucleoprotein complexes (RNP). Approximately 100 µL of cells from freshly grown parental strains on TAP plates were resuspended in 1 mL TAP medium and centrifuged at 600 × g for 2 min. The cell pellet was resuspended in 100 µL TAP medium, 100 µL autolysin and incubated for 15 min at 20 °C to remove cell walls. Cells were then incubated at 40 °C for 30 min with gentle agitation, washed three times with 1 mL TAPS (TAP supplemented with 40 mM sucrose) and finally resuspended in approximately 80 µL TAPS per 10 µL cell pellet. For each transfection, 40 µL of cell suspension was mixed with 4 µL RNP and 1.2 µL linearized donor template DNA, electroporated in 10 µL replicates using the Neon electroporation system at 2300 V for 13 ms and 3 pulses (Thermo Fisher Scientific, #MPK5000) and recovered in 1 mL TAPS overnight. The following day, cells were pelleted and spread on TAP plates with appropriate antibiotic selection (50 µg/mL blasticidin S (Invitrogen, #MSPP-ANT-BL-05) or 100 µg/mL spectinomycin (Chem-Impex, #14311)) and incubated in constant light until colonies formed (2-5 days). Successful knock-ins were confirmed by PCR amplification and sequencing.

### Total Internal Reflection Fluorescence (TIRF) microscopy

Prior to TIRF microscopy, 1 mL of M medium (1.7 mM NaCitrate, 37 µM FeCl_3_, 360 µM CaCl_2_, 1.2 mM MgSO_4_, 3.75 mM NH_4_NO_3_, 0.51 mM KH_2_PO_4_, 0.66 mM K_2_HPO_4_, 16 µM H_3_BO_3_, 3.5 µM ZnSO_4_, 2 µM MnSO_4_, 0.84 µM CoCl_2_, 0.83 µM Na_2_MoO_4_, 0.44 µM CuSO_4_, pH 6.8) was inoculated with *C. reinhardtii* cells and incubated shaking at 23 °C and 120 rpm with 14:10 h light:dark cycle for 2-3 days. Equal volumes of cell suspension and cr-TIRF buffer (5 mM HEPES pH 7.5, 5 mM EGTA pH 8.0) were mixed on glass coverslips (VWR, #48366-227), then mounted on glass microscope slides (Kemtech #0303-2122) using white petroleum (Vaseline, CAS #8009-03-8) and imaged at 23 °C. TIRF microscopy was performed using a Nikon Ti-E motorized inverted microscope with TIRF illuminator equipped with Agilent laser launch 488 and 561 nm laser lines and a Hamamatsu ImageEM EM-CCD camera fitted with DualView for simultaneous 2-color acquisition and 100x oil-immersion objective.

Kymographs were created from time lapse recordings in Fiji (v2.14.0) (Schindelin *et al*., 2012). Histogram matching bleach correction was performed for the 561 nm channel (mScarlet-IFT54). Spline-fitted segmented lines were placed along the flagellum’s length to reslice the stack into time-distance kymographs. Brightness and contrast were manually adjusted for each channel and image quality was enhanced using a Gaussian blur (sigma=0.5) followed by an unsharp mask (radius=1, mask=0.60). Individual protein tracks were traced by manual line selection and velocities calculated from distances covered over time (0.107 µm/pixel, 0.120 s/frame). Statistical analyses were conducted using scipy (v1.13.1) and statsmodels (v0.14.4) python libraries. Tukey’s HSD test was performed to assess differences in velocities across different channels and track directions. *P*-values were computed from the absolute difference between group means, dividing by the standard error of the differences and using the survival function of the studentized range distribution. All statistical tests were conducted with a significance level of α = 0.05.

### Protein sequence alignments

Protein sequences were retrieved from UniProt (https://www.uniprot.org/) for Homo sapiens, Sus scrofa and Mus musculus, from Phytozome (https://phytozome-next.jgi.doe.gov/) for C. reinhardtii, and TriTrypDB (https://tritrypdb.org/) (Aslett et al., 2010) for L. tarentolae. Protein sequence alignments of CFAP36 were generated using MAFFT in Snapgene (v8.0.2) and secondary structures of the human CFAP36 AlphaFold2 prediction (ID Q96G28) mapped with ENDscript2 (http://endscript.ibcp.fr) (Robert & Gouet, 2014).

### Quantification and statistical analysis

Proteomics experiments were performed with at least three replicates. For heatmaps, the scaled data was log2-transformed and a one-way ANOVA performed for each protein across experimental groups, followed by Benjamini-Hochberg FDR correction for multiple testing. Proteins with q-values below 0.01 were considered statistically significant, and those exhibiting a fold-change greater than 2 compared to the control group were retained. For volcano plots, the unscaled intensities were log2(x+1) transformed and comparison of experimental groups was conducted using pairwise Student’s t-tests. The resulting p-values were adjusted for multiple testing using Benjamini-Hochberg FDR correction and log2 fold changes were calculated as the difference between mean group values of the transformed data. Dashed lines in volcano plots indicate significance thresholds of p-value 0.05 and Fold Change of 2. Thresholds of cSPOC rankings were determined by the elbow point in the distribution of scores, detected using the KneeLocator algorithm of the kneed (v0.8.5) python library. Differences in immunofluorescence signal intensities and RT-qPCR were assessed using Tukey’s HSD test with a significance level of α = 0.05. Numbers of samples are indicated in text and in figure legends.

## Supplementary Information

**Figure S1.**
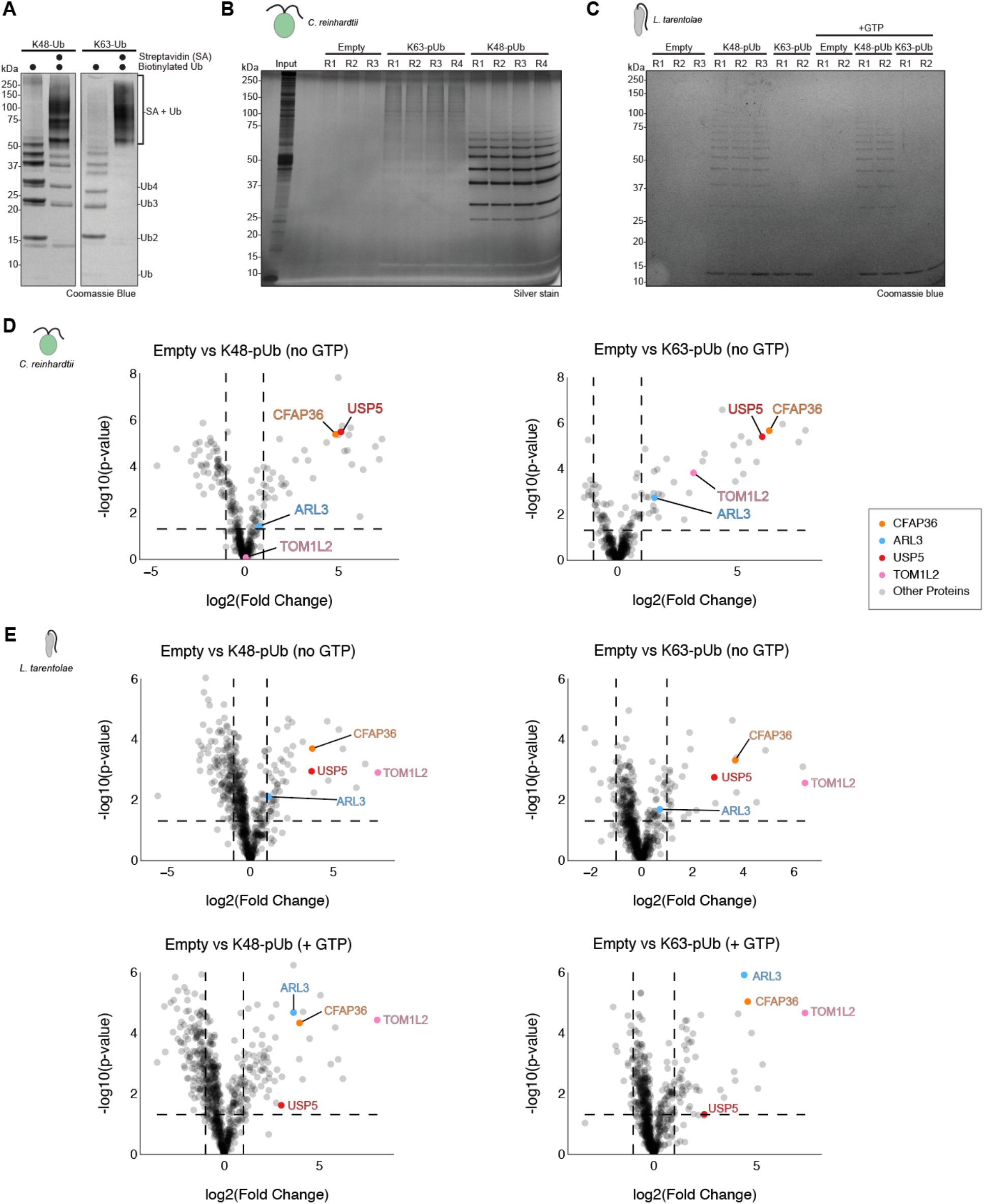
Proteomic identification of K48- and K63-linked ubiquitin readers in *Chlamydomonas reinhardtii* and *Leishmania tarentolae* flagella. Related to Figure 1. (A)SDS-PAGE analysis of biotinylated K48- and K63-linked polyubiquitin (pUb) chains before and after addition of streptavidin (SA) beads. (B-C) SDS-PAGE analysis of elution from empty and K48- or K63-linked ubiquitin-coated beads incubated with flagellar extracts from *C. reinhardtii* (B)and *L. tarentolae* (C). Each elution was analyzed by quantitative mass spectrometry. Replicate number is shown above each lane. (D)Volcano plots generated from TMT-MS analysis of flagellar extracts from *C. reinhardtii* comparing protein binding to empty beads versus beads coated with K48-linked (left) or K63-linked (right) polyubiquitin chains. Each plot shows significance versus enrichment. Individual proteins are shown as gray dots except for ARL3, CFAP36, TOM1L2, and USP5, which are colored and labeled. (E)Volcano plots generated from TMT-MS analysis of flagellar extracts from *L. tarentolae* comparing protein binding to empty beads versus beads coated with K48-linked (left) or K63-linked (right) ubiquitin chains in the absence (top) or presence (bottom) of GTP.

**Figure S2.**
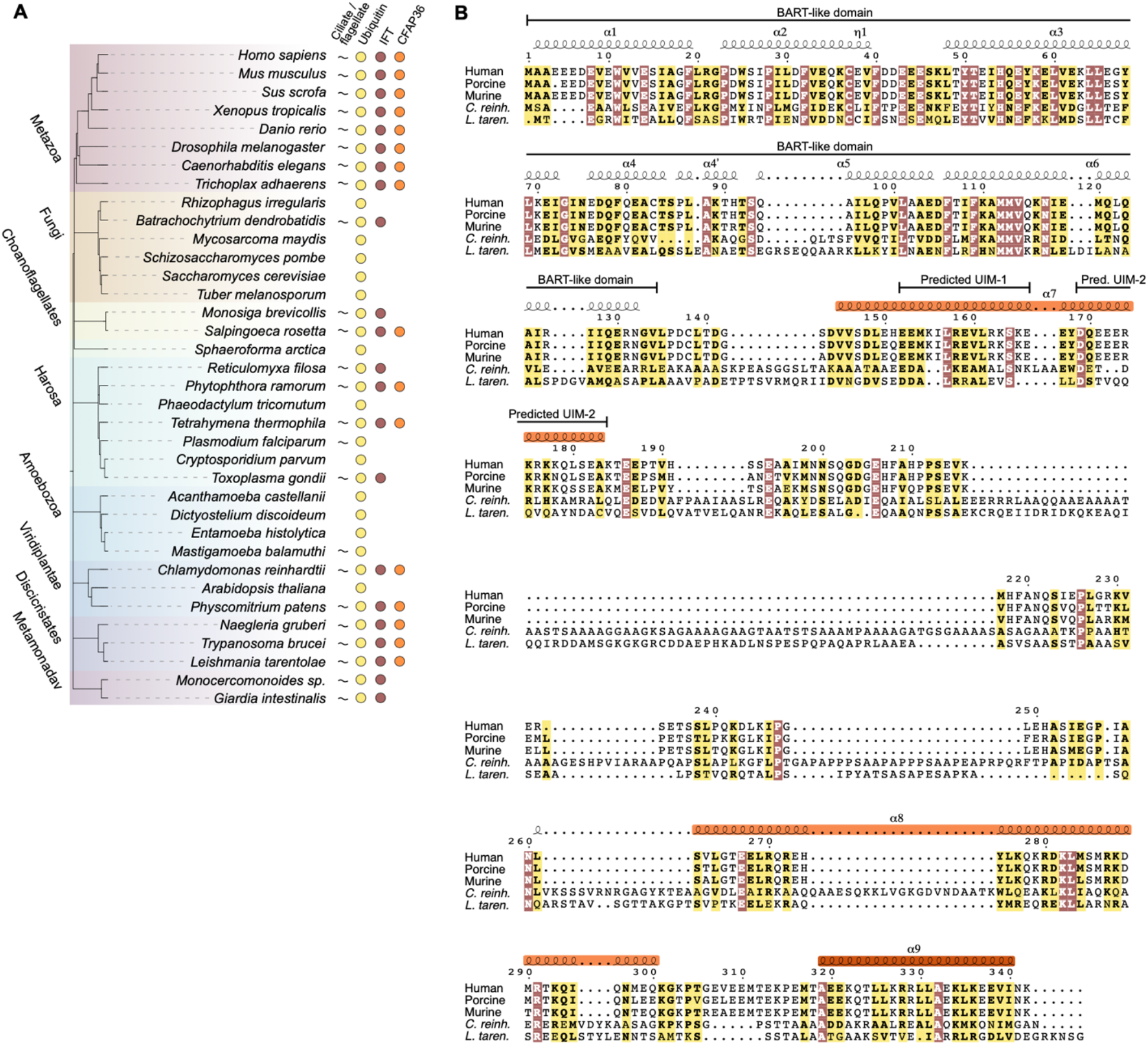
Phylogenetic and sequence conservation of CFAP36. Related to Figure 1. (A)A simplified phylogenetic tree illustrating the evolutionary relationships among various species, with indicators for the presence of cilia/flagella, ubiquitin, intraflagellar transport (IFT) proteins, and CFAP36. CFAP36 is only present in organisms with cilia, ubiquitin and IFT. (B)A multiple sequence alignment of CFAP36 protein sequences from the five organisms used in this study: Homo sapiens (human), Sus scrofa (porcine), Mus musculus (murine), Chlamydomonas reinhardtii (green algae), and Leishmania tarentolae (protozoan parasite). Secondary structure elements predicted from the human CFAP36 sequence are annotated above the alignment, with α-helices (α) indicated. The BART-like domain and predicted ubiquitin interacting motifs (UIMs) are labeled. Invariant residues, conserved across all five species, are highlighted in rust. Similar residues, sharing conserved properties, are highlighted in yellow.

**Figure S3.**
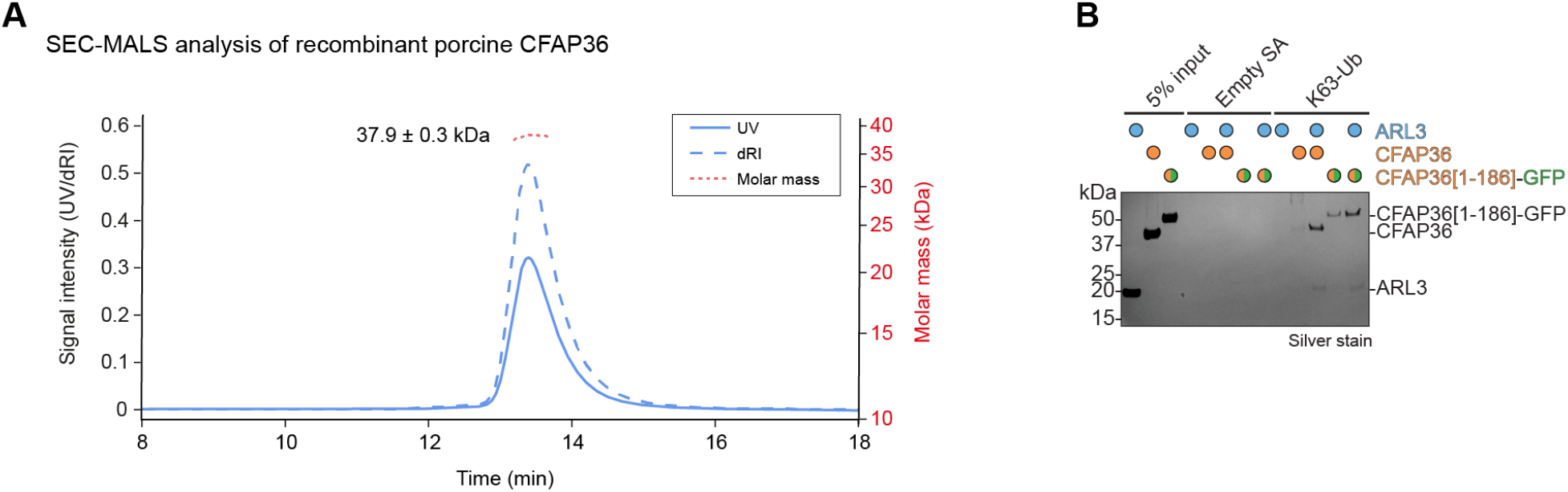
CFAP36 binds K48- and K63-linked polyubiquitin chains. Related to Figure 2. (A)Size exclusion chromatography coupled with multi-angle light scattering (SEC-MALS) of recombinant porcine CFAP36. The SEC elution profile is shown, with UV absorbance (blue), differential refractive index (dRI, dashed blue), and calculated molar mass (dashed red) plotted against time. The molar mass trace indicates a single species with a calculated molecular mass of 37.9 ± 0.3 kDa, consistent with a CFAP36 monomer. (B)Silver-stained SDS-PAGE analysis of a pulldown assay examining the binding interactions of recombinant full-length and truncated CFAP36 (residues 1-186 fused to a C-terminal GFP) with K63-linked ubiquitin (K63-Ub) in the presence and absence of ARL3. K63-Ub shows greater binding to truncated CFAP36 than full-length CFAP36, with ARL3 further enhancing the interaction.

**Figure S4.**
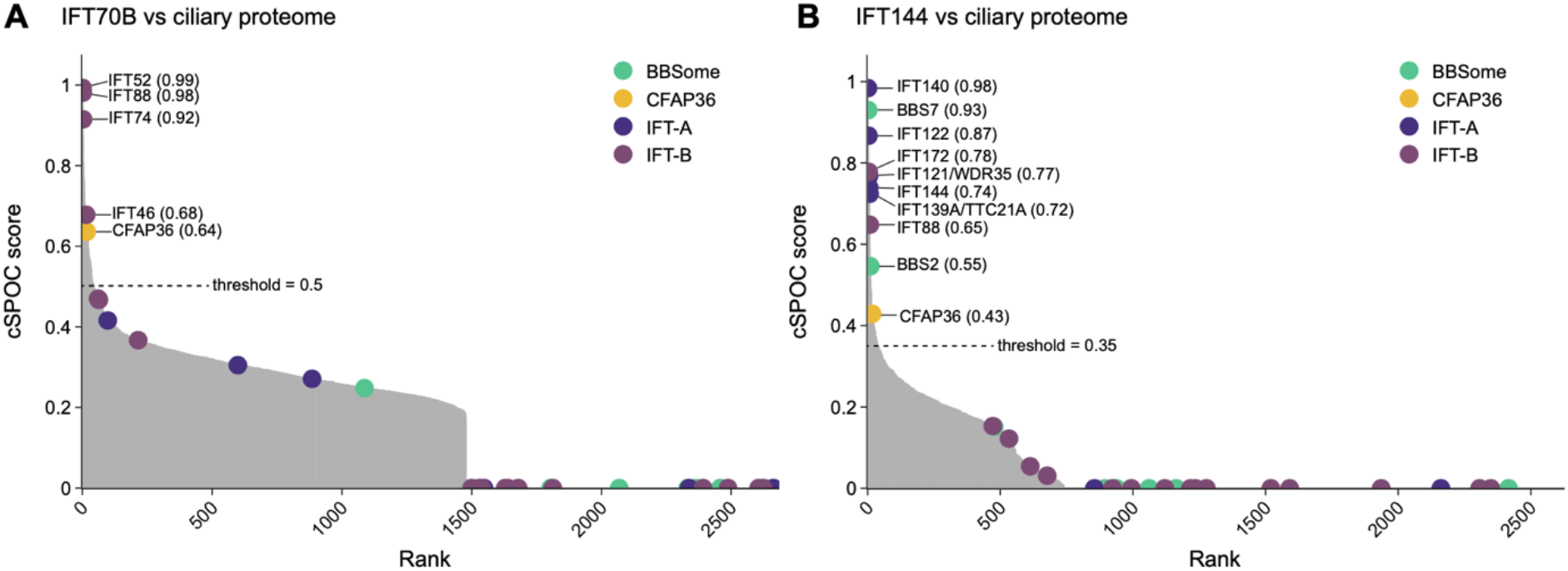
Identification of CFAP36 as an interacting partner of IFT70B and IFT144 in AlphaFold2-based screens. Related to Figure 5. (A-B) Plots illustrating cSPOC(Schmid & Walter, 2025) scores versus rank for pairwise AlphaFold2 predictions between a human ciliary proteome and IFT70B (A) and IFT144 (B). In each panel, the threshold for identifying a true interaction is determined by the elbow point of the curve. Pairwise predictions involving CFAP36 and subunits of the IFT-A, IFT-B and BBSome complexes are represented as colored dots. Those above the threshold are annotated with their protein name and cSPOC score.

**Figure S5.**
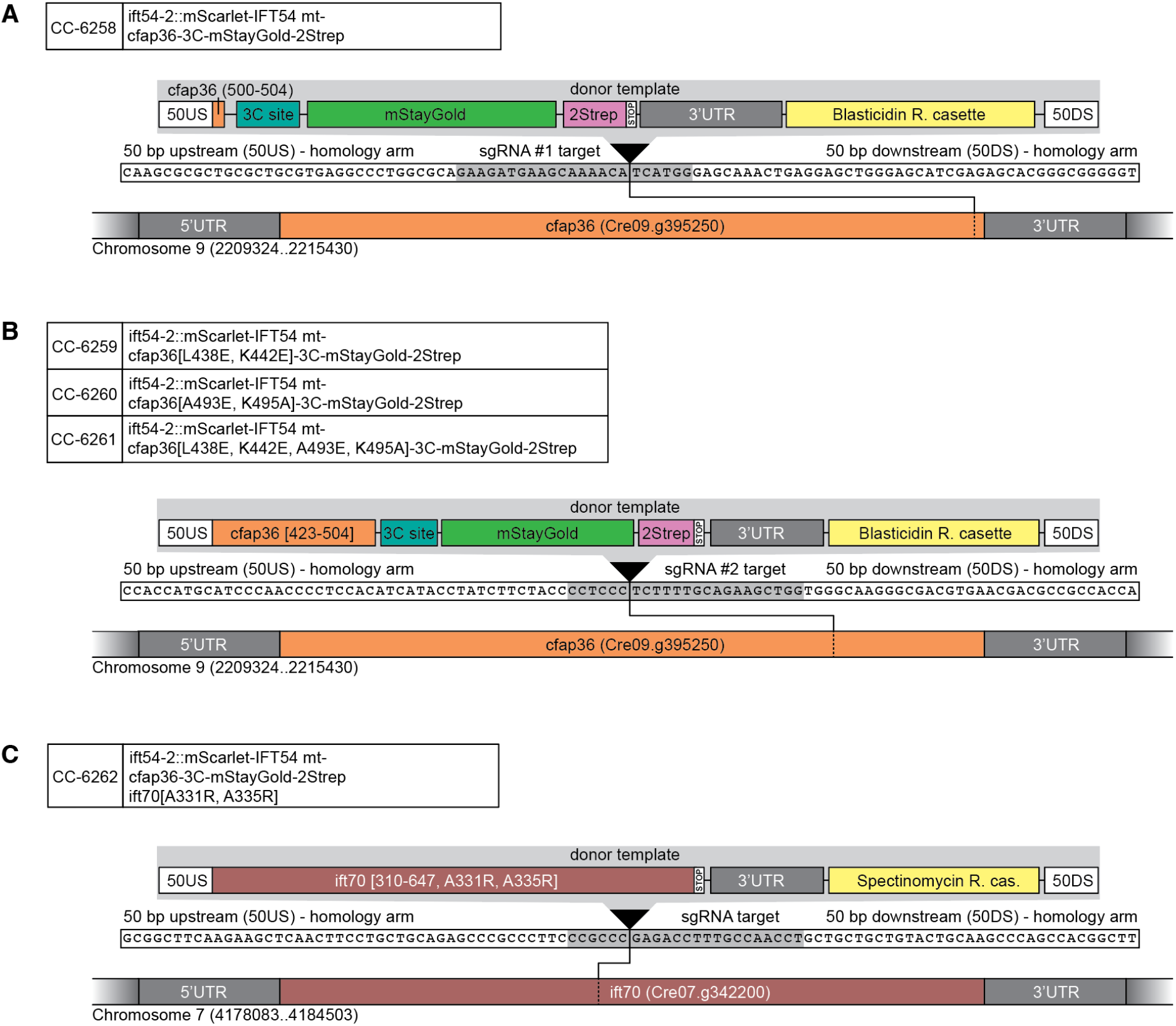
CRISPR/Cas9 donor templates. Related to Figures 5 and 7. (A-C) Schematics of CRISPR/Cas9 experimental design including sequence information of target sites used to (A) append an mStayGold fluorescent tag onto the C-terminus of CFAP36, (B) introduce point mutations into CFAP36 while also adding a C-terminal mStayGold fluorescent tag, and (C) to introduce point mutations into IFT70.

**Figure S6.**
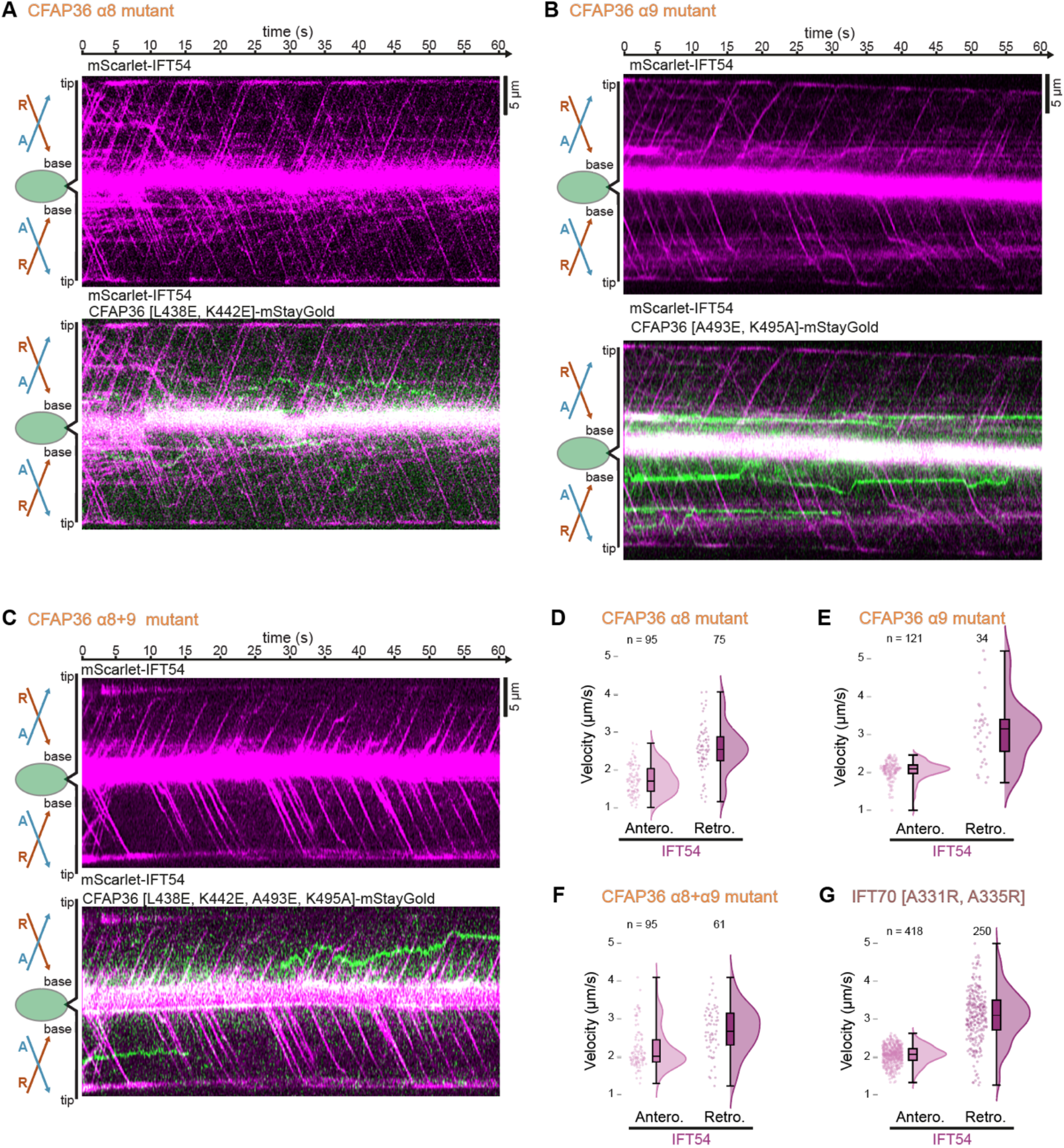
Retrograde transport of CFAP36 is abolished by mutations in predicted interfacial residues. Related to Figure 7. (A)Representative kymographs showing the movement of mScarlet-IFT54 alone (top) and overlaid with the CFAP36-mStayGold α8 [L438E, K442E] mutant (bottom) in *C. reinhardtii* flagella. (B)Representative kymographs showing the movement of mScarlet-IFT54 alone (top) and overlaid with the CFAP36-mStayGold α9 [A493E, K495A] mutant (bottom) in *C. reinhardtii* flagella. (C)Representative kymographs showing the movement of mScarlet-IFT54 alone (top) and overlaid with the CFAP36-mStayGold α8+α9 [L438E, K442E, A493E, and K495A] mutant (bottom) in *C. reinhardtii* flagella. The strong central signal intensities in panels A-C are due to epi-fluorescence signal from the cell body. (D-G) Raincloud plots showing quantification of anterograde and retrograde train velocities measured from mScarlet-IFT54 kymographs in strains expressing (D) CFAP36 α8 mutant, (E) CFAP36 α9 mutant, (F) CFAP36 α8+α9 mutant, and (G) IFT70 A331R, A335R mutant. Each raincloud plot shows individual data points, the density of each velocity range (represented by the cloud), and a boxplot. In the box plot, the central line represents the median, with the bottom and top of the box representing the first (Q1) and third (Q3) quartiles, respectively. The whiskers extend to the highest and the lowest measurements.

## Notes

### Competing Interest Statement

The authors have declared no competing interest.

## References

Alkanderi, S., Molinari, E., Shaheen, R., Elmaghloob, Y., Stephen, L. A., Sammut, V., Ramsbottom, S. A., Srivastava, S., Cairns, G., Edwards, N., Rice, S. J., Ewida, N., Alhashem, A., White, K., Miles, C. G., Steel, D. H., Alkuraya, F. S., Ismail, S. & Sayer, J. A. (2018). Am. J. Hum. Genet. 103, 612–620.

Allan, C., Burel, J.-M., Moore, J., Blackburn, C., Linkert, M., Loynton, S., Mac-Donald, D., Moore, W. J., Neves, C., Patterson, A., Porter, M., Tarkowska, A., Loranger, B., Avondo, J., Lagerstedt, I., Lianas, L., Leo, S., Hands, K., Hay, R. T., Patwardhan, A., Best, C., Kleywegt, G. J., Zanetti, G. & Swedlow, J. R. (2012). Nat. Methods 9, 245–253.

Argenzio, E., Bange, T., Oldrini, B., Bianchi, F., Peesari, R., Mari, S., Fiore, P. P. D., Mann, M. & Polo, S. (2011). Mol. Syst. Biol. 7, MSB2010118.

Aslett, M., Aurrecoechea, C., Berriman, M., Brestelli, J., Brunk, B. P., Carrington, M., Depledge, D. P., Fischer, S., Gajria, B., Gao, X., Gardner, M. J., Gingle, A., Grant, G., Harb, O. S., Heiges, M., Hertz-Fowler, C., Houston, R., Innamorato, F., Iodice, J., Kissinger, J. C., Kraemer, E., Li, W., Logan, F. J., Miller, J. A., Mitra, S., Myler, P. J., Nayak, V., Pennington, C., Phan, I., Pinney, D. F., Ramasamy, G., Rogers, M. B., Roos, D. S., Ross, C., Sivam, D., Smith, D. F., Srinivasamoorthy, G., Stoeckert, C. J., Subramanian, S., Thibodeau, R., Tivey, A., Treatman, C., Velarde, G. & Wang, H. (2010). Nucleic Acids Res 38, D457–D462.

Beneke, T., Demay, F., Wheeler, R. J. & Gluenz, E. (2020). Methods Mol. Biol. 2116, 485–495.

Christensen, S. T., Morthorst, S. K., Mogensen, J. B. & Pedersen, L. B. (2017). Cold Spring Harb. Perspect. Biol. 9, a028167.

Consortium, T. U., Bateman, A., Martin, M.-J., Orchard, S., Magrane, M., Ahmad, S., Alpi, E., Bowler-Barnett, E. H., Britto, R., Bye-A-Jee, H., Cukura, A., Denny, P., Dogan, T., Ebenezer, T., Fan, J., Garmiri, P., Gonzales, L. J. da C., Hatton-Ellis, E., Hussein, A., Ignatchenko, A., Insana, G., Ishtiaq, R., Joshi, V., Jyothi, D., Kandasaamy, S., Lock, A., Luciani, A., Lugaric, M., Luo, J., Lussi, Y., MacDougall, A., Madeira, F., Mahmoudy, M., Mishra, A., Moulang, K., Nightingale, A., Pundir, S., Qi, G., Raj, S., Raposo, P., Rice, D. L., Saidi, R., Santos, R., Speretta, E., Stephenson, J., Totoo, P., Turner, E., Tyagi, N., Vasudev, P., Warner, K., Watkins, X., Zaru, R., Zellner, H., Bridge, A. J., Aimo, L., Argoud-Puy, G., Auchincloss, A. H., Axelsen, K. B., Bansal, P., Baratin, D., Neto, T. M. B., Blatter, M.-C., Bolleman, J. T., Boutet, E., Breuza, L., Gil, B. C., Casals-Casas, C., Echioukh, K. C., Coudert, E., Cuche, B., Castro E. de, Estreicher, A., Famiglietti, M. L., Feuermann, M., Gasteiger, E., Gaudet, P., Gehant, S., Gerritsen, V., Gos, A., Gruaz, N., Hulo, C., Hyka-Nouspikel, N., Jungo, F., Kerhornou, A., Mercier, P. L., Lieberherr, D., Masson, P., Morgat, A., Muthukrishnan, V., Paesano, S., Pedruzzi, I., Pilbout, S., Pourcel, L., Poux, S., Pozzato, M., Pruess, M., Redaschi, N., Rivoire, C., Sigrist, C. J. A., Sonesson, K., Sundaram, S., Wu, C. H., Arighi, C. N., Arminski, L., Chen, C., Chen, Y., Huang, H., Laiho, K., McGarvey, P., Natale, D. A., Ross, K., Vinayaka, C. R., Wang, Q., Wang, Y. & Zhang, J. (2022). Nucleic Acids Res. 51, D523–D531.

Craig, R. J., Gallaher, S. D., Shu, S., Salomé, P. A., Jenkins, J. W., Blaby-Haas, C. E., Purvine, S. O., O’Donnell, S., Barry, K., Grimwood, J., Strenkert, D., Kropat, J., Daum, C., Yoshinaga, Y., Goodstein, D. M., Vallon, O., Schmutz, J. & Merchant, S. S. (2022). Plant Cell 35, 644– 672.

Craige, B., Brown, J. M. & Witman, G. B. (2013). Curr. Protoc. Cell Biol. 59, 3.41.1-3.41.9.

Crooks, G. E., Hon, G.Chandonia, J.-M. & Brenner, S. E. (2004). Genome Res. 14, 1188–1190.

Daggubati, V., Vykunta, A., Choudhury, A., Qadeer, Z., Mirchia, K., Saulnier, O., Zakimi, N., Hines, K., Paul, M., Wang, L., Jura, N., Xu, L., Reiter, J., Taylor, M., Weiss, W. & Raleigh, D. (2023). Res. Sq. rs.3.rs-3058335.

Desai, P. B., Stuck, M. W., Lv, B. & Pazour, G. J. (2020). J. Cell Biol. 219, e201912104.

Domire, J. S., Green, J. A., Lee, K. G., Johnson, A. D., Askwith, C. C. & Mykytyn, K. (2011). Cell. Mol. Life Sci. 68, 2951–2960.

Du, Y., Chen, F., Zhang, J., Lin, Z., Ma, Q., Xu, G., Xiao, D., Gui, Y., Yang, J. & Wan, S. (2019). Bone 127, 503–509.

Eguether, T., Agustin, J. T. S., Keady, B. T., Jonassen, J. A., Liang, Y., Francis, R., Tobita, K., Johnson, C. A., Abdelhamed, Z. A., Lo, C. W. & Pazour, G. J. (2014). Dev. Cell 31, 279–290.

Eng, J. K., Jahan, T. A. & Hoopmann, M. R. (2013). PROTEOMICS 13, 22– 24.

Engel, B. D., Lechtreck, K.-F., Sakai, T., Ikebe, M., Witman, G. B. & Marshall, W. F. (2009). Methods Cell Biol. 93, 157–177.

Evans, R. J., Schwarz, N., Nagel-Wolfrum, K., Wolfrum, U., Hardcastle, A. J. & Cheetham, M. E. (2010). Hum. Mol. Genet. 19, 1358–1367.

Evans, R., O’Neill, M., Pritzel, A., Antropova, N., Senior, A., Green, T., Žídek, A., Bates, R., Blackwell, S., Yim, J., Ronneberger, O., Bodenstein, S., Zielinski, M., Bridgland, A., Potapenko, A., Cowie, A., Tunyasuvunakool, K., Jain, R., Clancy, E., Kohli, P., Jumper, J. & Hassabis, D. (2021). BioRxiv 2021.10.04.463034.

Galcheva-Gargova, Z., Theroux, S. J. & Davis, R. J. (1995). Oncogene 11, 2649–2655.

Gui, M., Farley, H., Anujan, P., Anderson, J. R., Maxwell, D. W., Whitchurch, J. B., Botsch, J. J., Qiu, T., Meleppattu, S., Singh, S. K., Zhang, Q., Thompson, J., Lucas, J. S., Bingle, C. D., Norris, D. P., Roy, S. & Brown, A. (2021). Cell 184, 5791-5806.e19.

Hansen, J. N., Rassmann, S., Stüven, B., Jurisch-Yaksi, N. & Wachten, D. (2021). Eur. Phys. J. E 44, 18.

Hofmann, K. & Falquet, L. (2001). Trends Biochem. Sci. 26, 347–350.

Holtan, J. P., Teigen, K., Aukrust, I., Bragadóttir, R. & Houge, G. (2019). Ophthalmic Genet. 40, 124–128.

Huang, F., Kirkpatrick, D., Jiang, X., Gygi, S. & Sorkin, A. (2006). Mol. Cell 21, 737–748.

Huang, K., Diener, D. R. & Rosenbaum, J. L. (2009). J. Cell Biol. 186, 601– 613.

Huang, Y., Dong, X., Sun, S. Y., Lim, T.-K., Lin, Q. & He, C. Y. (2024). Sci. Adv. 10, eadq2950.

Ivorra-Molla, E., Akhuli, D., McAndrew, M. B. L., Scott, W., Kumar, L., Palani, S., Mishima, M., Crow, A. & Balasubramanian, M. K. (2024). Nat. Biotechnol. 42, 1368–1371.

Jin, H., White, S. R., Shida, T., Schulz, S., Aguiar, M., Gygi, S. P., Bazan, J. F. & Nachury, M. V. (2010). Cell 141, 1208–1219.

Jumper, J., Evans, R., Pritzel, A., Green, T., Figurnov, M., Ronneberger, O., Tunyasuvunakool, K., Bates, R., Žídek, A., Potapenko, A., Bridgland, A., Meyer, C., Kohl, S. A. A., Ballard, A. J., Cowie, A., Romera-Paredes, B., Nikolov, S., Jain, R., Adler, J., Back, T., Petersen, S., Reiman, D., Clancy, E., Zielinski, M., Steinegger, M., Pacholska, M., Berghammer, T., Bodenstein, S., Silver, D., Vinyals, O., Senior, A. W., Kavukcuoglu, K., Kohli, P. & Hassabis, D. (2021). Nature 596, 583–589.

Keady, B. T., Samtani, R., Tobita, K., Tsuchya, M., Agustin, J. T. S., Follit, J. A., Jonassen, J. A., Subramanian, R., Lo, C. W. & Pazour, G. J. (2011). Dev. Cell 22, 940–951.

Labun, K., Montague, T. G., Krause, M., Cleuren, Y. N. T., Tjeldnes, H. & Valen, E. (2019). Nucleic Acids Res. 47, W171–W174.

Lacey, S. E., Foster, H. E. & Pigino, G. (2023). Nat. Struct. Mol. Biol. 30, 584– 593.

Lacey, S. E., Graziadei, A. & Pigino, G. (2024). Cell 187, 4621-4636.e18.

Lauer, J., Segeletz, S., Cezanne, A., Guaitoli, G., Raimondi, F., Gentzel, M., Alva, V., Habeck, M., Kalaidzidis, Y., Ueffing, M., Lupas, A. N., Gloeckner, C. J. & Zerial, M. (2019). ELife 8, e46302.

Lechtreck, K. (2022). J Cell Sci 135, 10.1242/jcs.260408.

Lechtreck, K. F., Brown, J. M., Sampaio, J. L., Craft, J. M., Shevchenko, A., Evans, J. E. & Witman, G. B. (2013). J. Cell Biol. 201, 249–261.

Lechtreck, K.-F., Johnson, E. C., Sakai, T., Cochran, D., Ballif, B. A., Rush, J., Pazour, G. J., Ikebe, M. & Witman, G. B. (2009). J. Cell Biol. 187, 1117–1132.

Li, B., Rauhauser, A. A., Dai, J., Sakthivel, R., Igarashi, P., Jetten, A. M. & Attanasio, M. (2011). Hum. Mol. Genet. 20, 4155–4166.

Liu, Y.-X., Sun, W.-Y., Xue, B., Zhang, R.-K., Li, W.-J., Xie, X. & Fan, Z.-C. (2022). J. Cell Biol. 221, e202111076.

Lokaj, M., Kösling, S. K., Koerner, C., Lange, S. M., Beersum, S. E. C. van, Reeuwijk, J. van, Roepman, R., Horn, N., Ueffing, M., Boldt, K. & Wittinghofer, A. (2015). Struct.(Lond., Engl.:1993) 23, 2122–2132.

Lv, B., Stuck, M. W., Desai, P. B., Cabrera, O. A. & Pazour, G. J. (2021). J. Cell Biol. 220, e202010177.

Mali, G., Issa, K. H. B., Ren, M., Burnet, B., Lu, H., Melia, C., Heesom, K. & Roy, S. (2024). 10.21203/rs.3.rs-5233849/v1.

May, E. A., Kalocsay, M., D’Auriac, I. G., Schuster, P. S., Gygi, S. P., Nachury, M. V. & Mick, D. U. (2021). J. Cell Biol. 220, e202007207.

McCafferty, C. L., Papoulas, O., Lee, C., Bui, K. H., Taylor, D. W., Marcotte, E. M. & Wallingford, J. B. (2024). Dev. Cell 10.1016/j.devcel.2024.11.019.

Meng, E. C., Goddard, T. D., Pettersen, E. F., Couch, G. S., Pearson, Z. J., Morris, J. H. & Ferrin, T. E. (2023). Protein Sci. 32, e4792.

Mercey, O., Mukherjee, S., Guichard, P. & Hamel, V. (2024). Curr. Opin. Cell Biol. 88, 102361.

Mick, D. U., Rodrigues, R. B., Leib, R. D., Adams, C. M., Chien, A. S., Gygi, S. P. & Nachury, M. V. (2015). Dev. Cell 35, 497–512.

Mirdita, M., Schütze, K., Moriwaki, Y., Heo, L., Ovchinnikov, S. & Steinegger, M. (2022). Nat. Methods 19, 679–682.

Mitchison, H. M. & Valente, E. M. (2017). J. Pathol. 241, 294–309.

Moran, A. L., Louzao-Martinez, L., Norris, D. P., Peters, D. J. M. & Blacque, O. E. (2024). Nat. Rev. Nephrol. 20, 83–100.

Mukhopadhyay, S., Wen, X., Chih, B., Nelson, C. D., Lane, W. S., Scales, S. J. & Jackson, P. K. (2010). Gene Dev 24, 2180–2193.

Mukhopadhyay, S., Wen, X., Ratti, N., Loktev, A., Rangell, L., Scales, S. J. & Jackson, P. K. (2013). Cell 152, 210–223.

Niehrs, C., Silva, F. D. & Seidl, C. (2025). Trends Cell Biol. 35, 24–32.

Nievergelt, A. P., Diener, D. R., Bogdanova, A., Brown, T. & Pigino, G. (2023). Cell Rep. Methods 3, 100562.

Obayashi, T., Kodate, S., Hibara, H., Kagaya, Y. & Kinoshita, K. (2022). Nucleic Acids Res. 51, D80–D87.

Ojeda-Naharros, I., Das, T., Castro, R. A., Bazan, J. F., Vaisse, C. & Nachury, M. V. (2025). PLOS Biol. 23, e3003025.

Pazour, G. J., Agrin, N., Leszyk, J. & Witman, G. B. (2005). J. Cell Biol. 170, 103–113.

Perez-Riverol, Y., Bandla, C., Kundu, D. J., Kamatchinathan, S., Bai, J., Hewapathirana, S., John, N. S., Prakash, A., Walzer, M., Wang, S. & Vizcaíno, J. A. (2024). Nucleic Acids Res. 53, D543–D553.

Pigino, G., Geimer, S., Lanzavecchia, S., Paccagnini, E., Cantele, F., Diener, D. R., Rosenbaum, J. L. & Lupetti, P. (2009). J. Cell Biol. 187, 135–148.

Pir, M. S., Begar, E., Yenisert, F., Demirci, H. C., Korkmaz, M. E., Karaman, A., Tsiropoulou, S., Firat-Karalar, E. N., Blacque, O. E., Oner, S. S., Doluca, O., Cevik, S. & Kaplan, O. I. (2024). Nucleic Acids Res. 52, 8127–8145.

Puertollano, R. (2005). J. Biol. Chem. 280, 9258–9264.

Reck-Peterson, S. L., Redwine, W. B., Vale, R. D. & Carter, A. P. (2018). Nat. Rev. Mol. Cell Biol. 19, 382–398.

Robert, X. & Gouet, P. (2014). Nucleic Acids Research 42, W320–4.

Rohatgi, R., Milenkovic, L. & Scott, M. P. (2007). Science 317, 372–376.

Sato, Y., Yoshikawa, A., Mimura, H., Yamashita, M., Yamagata, A. & Fukai, S. (2009). EMBO J. 28, 2461–2468.

Satopää, V., Albrecht, J., Irwin, D. & Raghavan, B. (2011). 2011 31st Int. Conf. Distrib. Comput. Syst. Work. 166–171.

Schindelin, J., Arganda-Carreras, I., Frise, E., Kaynig, V., Longair, M., Pietzsch, T., Preibisch, S., Rueden, C., Saalfeld, S., Schmid, B., Tinevez, J.-Y., White, D. J., Hartenstein, V., Eliceiri, K., Tomancak, P. & Cardona, A. (2012). Nat. Methods 9, 676–682.

Schmid, E. W. & Walter, J. C. (2025). Mol. Cell 10.1016/j.molcel.2025.01.034.

Schweke, H., Pacesa, M., Levin, T., Goverde, C. A., Kumar, P., Duhoo, Y., Dornfeld, L. J., Dubreuil, B., Georgeon, S., Ovchinnikov, S., Woolfson, D. N., Correia, B. E., Dey, S. & Levy, E. D. (2024). Cell 187, 999-1010.e15.

Seo, S., Zhang, Q., Bugge, K., Breslow, D. K., Searby, C. C., Nachury, M. V. & Sheffield, V. C. (2011). PLoS Genet. 7, e1002358.

Shiba, Y., Katoh, Y., Shiba, T., Yoshino, K., Takatsu, H., Kobayashi, H., Shin, H.-W., Wakatsuki, S. & Nakayama, K. (2004). J. Biol. Chem. 279, 7105– 7111.

Shinde, S. R., Mick, D. U., Aoki, E., Rodrigues, R. B., Gygi, S. P. & Nachury, M. V. (2023). Dev. Cell 58, 677-693.e9.

Shinde, S. R., Nager, A. R. & Nachury, M. V. (2020). J. Cell Biol. 219, 199.

Sigrist, C. J. A., Castro E. de, Cerutti, L., Cuche, B. A., Hulo, N., Bridge, A., Bougueleret, L. & Xenarios, I. (2013). Nucleic Acids Res. 41, D344– D347.

Strom, S. P., Clark, M. J., Martinez, A., Garcia, S., Abelazeem, A. A., Matynia, A., Parikh, S., Sullivan, L. S., Bowne, S. J., Daiger, S. P. & Gorin, M. B. (2016). PLoS ONE 11, e0150944.

Swatek, K. N., Usher, J. L., Kueck, A. F., Gladkova, C., Mevissen, T. E. T., Pruneda, J. N., Skern, T. & Komander, D. (2019). Nature 572, 533–537.

Szklarczyk, D., Kirsch, R., Koutrouli, M., Nastou, K., Mehryary, F., Hachilif, R., Gable, A. L., Fang, T., Doncheva, N. T., Pyysalo, S., Bork, P., Jensen, L. J. & Mering, C. von (2022). Nucleic Acids Res. 51, D638–D646.

Taipale, J., Cooper, M. K., Maiti, T. & Beachy, P. A. (2002). Nature 418, 892–897.

Tubiana, J., Schneidman-Duhovny, D. & Wolfson, H. J. (2022). Nat. Methods 19, 730–739.

Uhlén, M., Fagerberg, L., Hallström, B. M., Lindskog, C., Oksvold, P., Mardinoglu, A., Sivertsson, Å., Kampf, C., Sjöstedt, E., Asplund, A., Olsson, I., Edlund, K., Lundberg, E., Navani, S., Szigyarto, C. A.-K., Odeberg, J., Djureinovic, D., Takanen, J. O., Hober, S., Alm, T., Edqvist, P.-H., Berling, H., Tegel, H., Mulder, J., Rockberg, J., Nilsson, P., Schwenk, J. M., Hamsten, M., Feilitzen, K. von, Forsberg, M., Persson, L., Johansson, F., Zwahlen, M., Heijne, G. von, Nielsen, J. & Pontén, F. (2015). Sci. (N. York, NY) 347, 1260419.

Varadi, M., Bertoni, D., Magana, P., Paramval, U., Pidruchna, I., Radhakrishnan, M., Tsenkov, M., Nair, S., Mirdita, M., Yeo, J., Kovalevskiy, O., Tunyasuvunakool, K., Laydon, A., Žídek, A., Tomlinson, H., Hariharan, D., Abrahamson, J., Green, T., Jumper, J., Birney, E., Steinegger, M., Hassabis, D. & Velankar, S. (2023). Nucleic Acids Res. 52, D368– D375.

Vasquez, S. S. V., Dam, J.van & Wheway, G. (2021). Mol. Biol. Cell 32, br13.

Veltel, S., Gasper, R., Eisenacher, E. & Wittinghofer, A. (2008). Nat. Struct. Mol. Biol. 15, 373–380.

Wang, J., Kidmose, R. T., Boegholm, N., Zacharia, N. K., Thomsen, M. B., Christensen, A., Malik, T., Lechtreck, K. & Lorentzen, E. (2025). J. Biol. Chem. 301, 108237.

Wilkinson, K. D., Tashayev, V. L., O’Connor, L. B., Larsen, C. N., Kasperek, E. & Pickart, C. M. (1995). Biochemistry 34, 14535–14546.

Wingfield, J. L., Mengoni, I., Bomberger, H. & Jiang, Y. Y. (2017). Elife.

Wright, K. J., Baye, L. M., Olivier-Mason, A., Mukhopadhyay, S., Sang, L., Kwong, M., Wang, W., Pretorius, P. R., Sheffield, V. C., Sengupta, P., Slusarski, D. C. & Jackson, P. K. (2011). Genes Dev. 25, 2347–2360.

Yildiz, O. & Khanna, H. (2012). Vis. Res. 75, 112–116.

Young, P., Deveraux, Q., Beal, R. E., Pickart, C. M. & Rechsteiner, M. (1998). J. Biol. Chem. 273, 5461–5467.

Zhang, Q., Nishimura, D., Seo, S., Vogel, T., Morgan, D. A., Searby, C., Bugge, K., Stone, E. M., Rahmouni, K. & Sheffield, V. C. (2011). Proc Natl Acad Sci USA 108, 20678–20683.

Zhang, Q., Nishimura, D., Vogel, T., Shao, J., Swiderski, R., Yin, T., Searby, C., Carter, C. S., Kim, G., Bugge, K., Stone, E. M. & Sheffield, V. C. (2013). J. Cell. Sci. 126, 2372–2380.

Zimmermann, L., Stephens, A., Nam, S.-Z., Rau, D., Kübler, J., Lozajic, M., Gabler, F., Söding, J., Lupas, A. N. & Alva, V. (2018). J. Mol. Biol. 430, 2237–2243.

